# Distributed functions of prefrontal and parietal cortices during sequential categorical decisions

**DOI:** 10.1101/2020.05.21.108910

**Authors:** Yang Zhou, Matthew Rosen, Sruthi K. Swaminathan, Nicolas Y. Masse, Oliver Zhu, David J. Freedman

## Abstract

The ability to compare sequential sensory inputs is crucial for solving many behavioral tasks. To understand the neuronal mechanisms underlying sequential decisions, we compared neuronal responses in the prefrontal cortex (PFC) and the lateral and medial intra-parietal (LIP and MIP) areas in monkeys trained to decide whether sequentially presented stimuli were from matching (M) or nonmatching (NM) categories. We found that PFC leads the M/NM decision process relying on nonlinear neuronal integration of sensory and mnemonic information, whereas LIP and MIP are more involved in sensory evaluation and motor planning, respectively. Furthermore, multi-module recurrent neural networks trained on the same task exhibited the key features of PFC and LIP encoding, including nonlinear integrative encoding in the PFC-like module which was crucial for M/NM decisions. Together, our results illuminate the relative functions of LIP, PFC, and MIP in sensory, cognitive and motor functions, and suggest that nonlinear integration of task-related variables in PFC is important for mediating sequential decisions.

## Introduction

The ability to compare and make decisions about sequentially presented sensory stimuli is essential for generating appropriate behavioral responses to the stimuli and events in our surroundings. Such sequential decisions require incoming sensory information to be compared to information maintained in short-term memory. Although previous work has given insight into patterns of activity across several cortical areas during sequential decision tasks, particularly those based on delayed match and delayed non-match to sample paradigms (Freedman and Assad, 2006; Freedman et al., 2001; Miller and Desimone, 1994; Miller et al., 1991; Wallis et al., 2001), the mechanisms and computations underlying sequential decisions remain largely unknown. The current study examines the pattern of neuronal activity across three interconnected frontal-parietal cortical areas—PFC, LIP, and MIP—during a delayed match to category (DMC) task which requires monkeys to indicate whether sequentially presented sample and test stimuli belong to the same category.

The lateral PFC, LIP and MIP are important processing stages for mediating decision-making tasks (de Lafuente et al., 2015; Ding and Gold, 2012; Gold and Shadlen, 2007; Huk et al., 2017; Kim and Shadlen, 1999; Padoa-Schioppa and Assad, 2006; Platt and Glimcher, 1999; Rossi-Pool et al., 2017; Shadlen and Newsome, 1996; Sugrue et al., 2004). PFC is involved in higher cognitive and executive functions, including short-term working memory (Funahashi et al., 1993; Miller et al., 1996; Romo et al., 1999), top-down attentional control (Gregoriou et al., 2009; Gregoriou et al., 2014; Moore and Armstrong, 2003; Schafer and Moore, 2011; Squire et al., 2013), response inhibition (Schall and Godlove, 2012; Stuphorn and Schall, 2006), learning (Antzoulatos and Miller, 2011; Asaad et al., 1998; Brincat and Miller, 2015; Seger and Miller, 2010), and performance monitoring (Hasegawa et al., 2000; Stuphorn et al., 2000). Meanwhile, LIP and MIP are often associated with visuospatial processing related to attention and movement planning (Andersen and Buneo, 2002; Andersen et al., 1997; Bisley and Goldberg, 2003, 2010; Colby and Goldberg, 1999; Cui and Andersen, 2007; Gnadt and Andersen, 1988; Gottlieb et al., 1998; Malhotra et al., 2009; Snyder et al., 1997; Zhou et al., 2016), as well as more cognitive functions such as perceptual decisions (Hanks et al., 2006; Huk and Shadlen, 2005; Kiani and Shadlen, 2009; Platt and Glimcher, 1999; Roitman and Shadlen, 2002; Shadlen and Newsome, 1996; Yang and Shadlen, 2007; Zhou and Freedman, 2019) and categorization (Fitzgerald et al., 2011; Freedman and Assad, 2006; Swaminathan and Freedman, 2012; Swaminathan et al., 2013). However, these three cortical areas have not been directly compared in the same animals performing the same tasks, thus their specific roles in the sensory, cognitive, and motor aspects of decisions have not been precisely delineated.

We recorded neural activity in PFC, LIP, and MIP while monkeys performed a visual-motion DMC task in which they reported, by releasing a manual lever, whether the category of a test stimulus matched the category of a previous sample. Our group showed previously that neural activity in PFC, LIP, and MIP all encode learned categories during this task (Freedman and Assad, 2006; Swaminathan and Freedman, 2012; Swaminathan et al., 2013), with evidence suggesting that LIP is more closely involved in categorization of visual motion compared to MIP and PFC. However, key questions remain about the mechanisms by which sample and test stimuli are compared in order to generate M/NM decisions. In this study, we focus on the test period of the DMC task, during which monkeys made their M/NM decisions. We found that test-period activity in all three cortical areas was correlated with monkeys’ M/NM decisions, but in different ways. M/NM selectivity in PFC appeared with a shorter latency than in LIP and MIP, suggesting a leading role for PFC in that decision process. Meanwhile, test-period activity in LIP showed the strongest categorical encoding of both the previously presented sample stimulus and the currently visible test stimulus. In contrast, MIP activity primarily reflected the monkeys’ arm and/or hand movements used to report their decisions. These results indicate that LIP and MIP are more closely involved in evaluating sensory stimuli and planning motor responses, respectively, whereas PFC is more specifically involved in the M/NM decision.

Individual neurons in PFC and LIP, but not MIP, encoded both the remembered sample and currently visible test categories simultaneously during the test period. However, compared to PFC, LIP sample and test category encoding was more linearly combined and not correlated in strength. In contrast, the strength of sample and test category encoding was positively correlated, and more nonlinearly combined in the PFC population. This is consistent with LIP evaluating sample and test stimuli in a bottom-up manner, while PFC nonlinearly integrates sample and test information to generate more explicit M/NM encoding. Interestingly, we found that PFC neurons which nonlinearly encoded sample and test stimuli were more closely involved in the monkeys’ M/NM decisions compared to other PFC neurons. These PFC neurons showed stronger encoding of important task variables, and their activity was more correlated with the monkeys’ M/NM decisions. These results suggest that nonlinear integrative encoding in PFC may play a preferential role in computing M/NM decisions in the DMC task.

Previous studies from our group and others have trained artificial recurrent neural networks (RNNs) on the same behavioral tasks used in experimental neurophysiological studies—an approach that has proven helpful in exploring putative circuit computations underlying cognitive tasks, generating predictions for analyses of experimental data, and even has potential for enhancing capabilities of RNNs (Engel et al., 2015; Masse et al., 2019; Yang et al., 2019). To further understand the roles of nonlinear integration of task variables during M/NM decisions, we trained multi-module RNNs to perform the DMC task and analyzed neural activity within each module of the hidden layer during task performance. The trained RNN exhibited a high level of behavioral task performance, and the population of units in the hidden layers showed similar patterns of activity and dynamics as in neural data from PPC and PFC. In particular, this included decision-correlated nonlinear encoding in the higher order module of sample and test information during the decision period of the task—similar to that observed in the neural recordings from PFC. Importantly, we causally tested the hypothesis that the RNN was reliant on this nonlinear encoding for decision computation by specifically silencing the activity of the RNN units which showed nonlinear encoding. This caused an impairment of task performance that was substantially greater than silencing an equivalent number of units which did not exhibit such nonlinear encoding—supporting the importance of nonlinear encoding of task-relevant information for decision making. Together, our results from both neurophysiological experiments in trained monkeys, and analysis of neural activity in trained RNNs, suggest that nonlinear integrative encoding, like that observed in PFC, functions as a key neuronal substrate for mediating sequential decisions.

## Results

### Task and behavioral performance

Two monkeys performed a DMC task, in which they needed to: 1) categorize the sample stimulus; 2) remember it during the delay period; 3) categorize the following test stimulus; 4) determine whether the test was a categorical match to the sample; and 5) report that decision by releasing or holding a manual lever (**Figure 1A**). Stimuli consisted of six random-dot motion directions that were grouped into two arbitrary learned categories, with three motion directions per category (**Figure 1B**). This corresponds to a total of 36 possible sample-test direction combinations and four sample-test category combinations. Both monkeys performed the DMC task with high accuracy (monkey A: 91%; monkey B: 97%), and greater than 80% for all 36 stimulus conditions during recordings from each of the three areas (**Figure 1C-D**, monkey A: PFC = 93%± 4.6%, LIP = 93%±4.8%, MIP = 88%±6.3%; monkey B: PFC = 98%± 1.3%, LIP = 96%±4.1%, MIP = 98%±2.5%). Slightly higher error rates were observed on match compared to non-match trials for all three data sets of both monkeys (Supplementary **table 1**). These results indicate that monkeys reliably based their M/NM decisions on the category membership of both sample and test stimuli.

**Figure 1.**
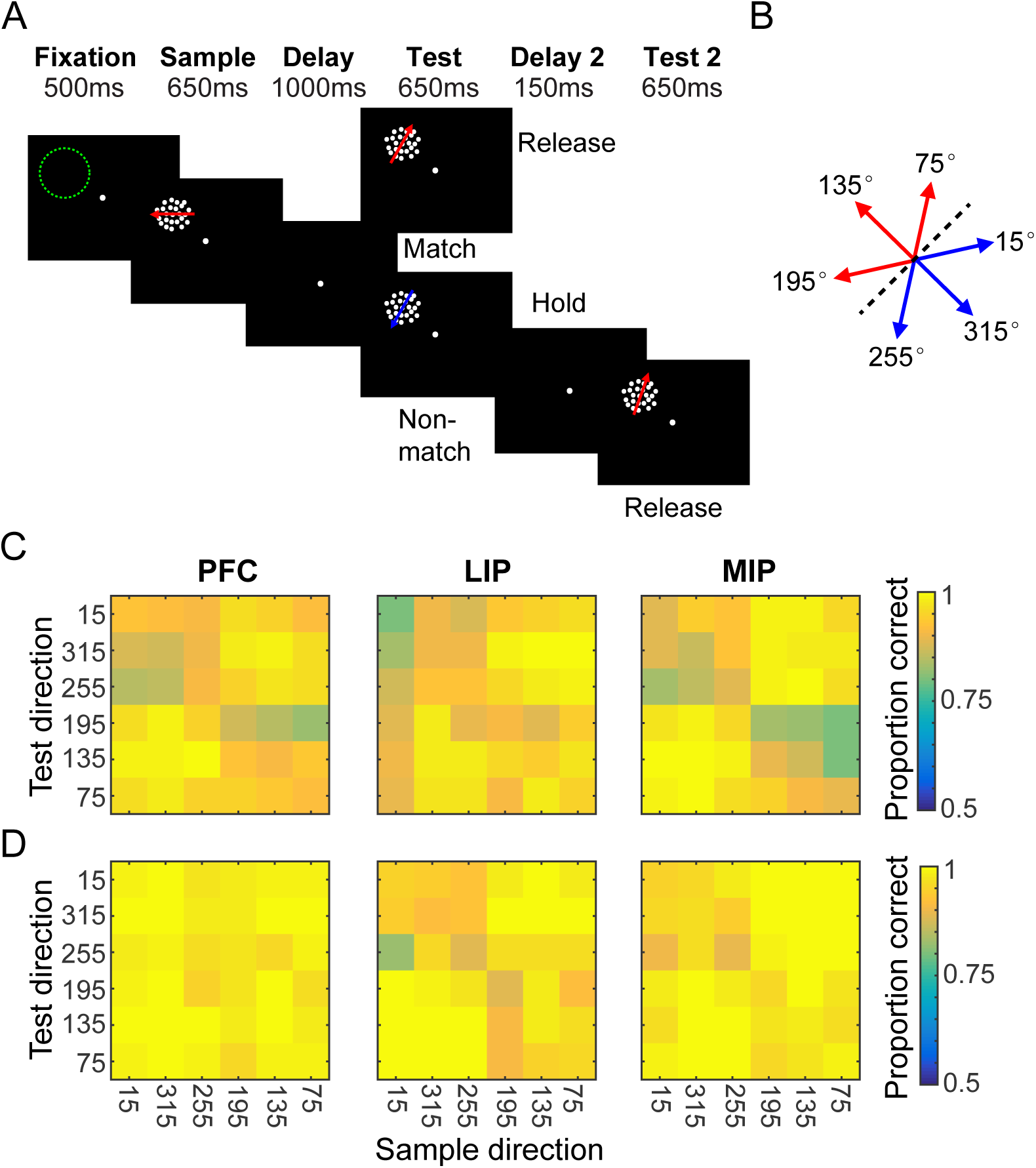
Task and behavior performance. **A**. Sequence of the DMC task. Monkeys needed to release a touch-bar when the categories of sample and test stimuli matched, or hold the bar and wait for the second test stimuli when they did not match. The M/NM decision was required to be made during the test period to receive the reward. Stimulus offset occurred immediately after monkeys released the touched-bar. The green dashed circle indicates the position of a neuron’s receptive field. **B**. Monkeys needed to group six motion directions into two categories (corresponding to the red and blue arrows) separated by a learned category boundary (black dashed line). **C-D**. Two monkeys’ average performance (accuracy) for all stimulus conditions during recordings from PFC, LIP and MIP recordings are shown separately. Each row corresponds to one monkey.

### M/NM selectivity in PFC, LIP, and MIP

We recorded neuronal spiking activity and local field potential (LFP) signals from PFC, LIP, and MIP in separate sessions; in each session, one cortical area was targeted while the monkey performed the DMC task (except for a subset of simultaneous PFC-LIP sessions). While neural data from the same sessions was presented in previous reports from our lab (Swaminathan and Freedman, 2012; Swaminathan et al., 2013), the current analysis focuses on decision signals in the test period of the task, which was not a primary focus of our previous work. Furthermore, our previous reports compared only pairs of cortical areas, rather than all three areas as in the current study. We focus our analysis on neurons which showed firing rates above an arbitrary threshold (the maximum of the condition averaged firing rate across the task period >= 5 sp/s), and were significantly modulated by task variables (see Materials and methods). In order to capture the neuronal correlates of the M/NM decision from sensory processing to motor planning, we analyzed neuronal activity from 50 to 350 ms following test stimulus onset, since monkeys’ manual responses (indicating their M/NM decisions) occurred within this epoch on 95.6% of match trials (monkey A: 246.2±14.0 ms, monkey B: 282.7±16.3 ms). Note that most of our subsequent analyses focus on a shorter-duration time window, ending prior to the monkeys mean reaction time. In this time period, a substantial fraction of neurons in each cortical area (PFC: 104/145, LIP: 35/53, MIP: 60/66) showed significantly different test-period activity between match and non-match trials (match vs. non-match; p < 0.01, one-way ANOVA). We classified these neurons as ‘match-preferring’ and ‘non-match-preferring’ based on whether they showed significantly higher firing rates for match or non-match trials, respectively (example neurons in **Figure 2-figure supplement 1** and population activity in **Figure 2A-C**).

**Figure 2.**
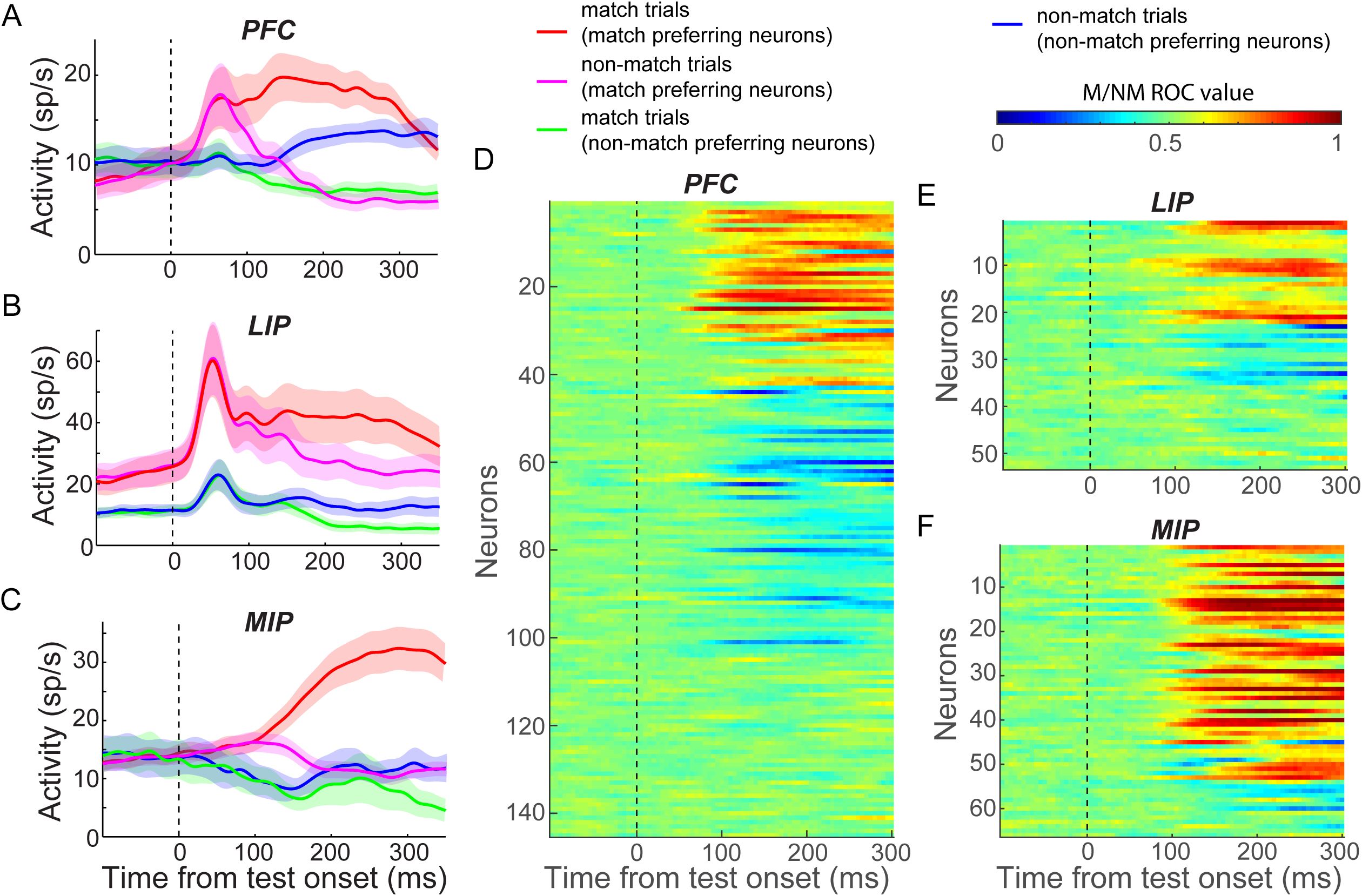
M/NM selectivity in PFC, LIP and MIP. **A-C**. The population activity of both match-preferring and non-match-preferring neurons in PFC (A), LIP (B) and MIP (C). The shaded area represents ±SEM. **D-F**. The strength of the M/NM selectivity was evaluated using ROC analysis for all neurons in PFC (D), LIP (E) and MIP (F). Values close to 0.0 and 1.0 correspond to strong encoding preference for non-match and match, respectively. Values of 0.5 indicate no M/NM selectivity.

To characterize the time-course of M/NM selectivity, we performed a receiver-operating characteristic (ROC) analysis comparing each neuron’s firing rates on match and non-match trials using a 50 ms window advanced in 5 ms steps (**Figure 2D-F**). Both match and non-match preferring neurons showed M/NM selectivity shortly after test onset in all three areas, though the fractions of neurons preferring match or non-match differed among the areas. PFC showed a relatively balanced distribution of match- and non-match-preferring neurons (match:non-match, 43:61), while LIP and MIP were more biased toward ‘match neurons’ (match:non-match, LIP: 22:13, MIP: 51:9, P_(PFCvs.LIP)_ = 0.027, P_(PFCvs.MIP)_ = 5.2×10^−8^, chi-square test). In all three areas, match-preferring neurons exhibited significantly greater activity than non-match-preferring neurons during the test period (P_(PFC)_ = 0.0372; P_(MIP)_ = 0.0427; P_(LIP)_ = 0.012; Wilcoxon test).

To compare the strength of M/NM selectivity between match and non-match preferring neurons, we calculated the unbiased fraction of explained variance (FEV) by the M/NM choice (see Materials and methods). On average, match-preferring neurons showed significantly greater M/NM selectivity in all three areas (P_(PFC)_ = 2.6×10^−4^; P_(MIP)_ = 0.024; P_(LIP)_ = 0.039; Wilcoxon test, **Figure 2-figure supplement 2**), as well as significantly shorter latency of M/NM selectivity than non-match preferring neurons in PFC (118.0 vs. 128.5 ms, P = 0.0071, Wilcoxon test; see Materials and methods). Differences of M/NM selectivity remained statistically significant in PFC and MIP (in both magnitude and latency), but showed only a non-significant trend in LIP, after equating the mean firing rates between groups of neurons (see Materials and methods, P_(PFC, magnitude)_ = 0.0015; P_(PFC, latency)_ = 0.0251; P_(LIP)_ = 0.12; P_(MIP)_ = 0.040; Wilcoxon test). This suggests that the stronger M/NM selectivity of match-preferring neurons observed in these areas is unlikely to be explained by the higher firing rates of match-preferring neurons.

### Comparing the roles of PFC, LIP, and MIP in M/NM decisions

To elucidate the relative contributions of PFC, LIP and MIP to M/NM decisions, we first compared the time-course of M/NM selectivity across areas. Since match-preferring neurons showed earlier and stronger M/NM selectivity than non-match-preferring neurons, and the distributions of the two groups of neurons were different among areas, we compared the M/NM selectivity of match-preferring and non-match preferring neurons separately in each cortical area. Using an unbiased FEV analysis, we found that match-preferring neurons exhibited significantly shorter-latency (see Materials and methods) M/NM selectivity in PFC than in LIP and MIP (**Figure 3A**, PFC: n=40, LIP: n=50, MIP: n=22; P_(PFC vs. LIP)_ = 0.026, P_(PFC vs. MIP)_ = 0.0058, Wilcoxon test). Non-match preferring neurons showed a similar trend (**Figure 3B**), although the difference was not significant—likely due to the small population of non-match preferring neurons in LIP and MIP (PFC: n=50, LIP: n=12, MIP: n=9; P_(PFC vs. LIP)_ = 0.11, P_(PFC vs. MIP)_ = 0.13, Wilcoxon test). We also examined the M/NM selectivity of the LFP signal, which likely reflects the activity and/or computations within the local network (Burns et al., 2010; Logothetis et al., 2001). The mean amplitude of the LFP in PFC also showed significantly shorter-latency M/NM selectivity than in LIP and MIP (**Figure 3C**, P_(PFC vs. LIP)_ = 8.9×10^−4^, P_(PFC vs. MIP)_ = 0.014, Wilcoxon test). We determined that the shorter-latency M/NM selectivity in PFC compared to PPC was not due to differences in latency on sessions which targeted each brain area, as we observed similar results in a different data set from another study (conducted in different monkeys) using the DMC task in our lab (Masse et al., 2017), in which neuronal activity was recorded simultaneously from PFC and PPC using a semi-chronic multielectrode approach (see Materials and methods). As shown in **Figure 3-figure supplement 1**, PFC neurons showed significantly shorter-latency M/NM selectivity than PPC neurons in that study (p = 0.0158, Wilcoxon test). Furthermore, the raw LFP amplitude, which was recorded simultaneously from the two areas, showed shorter-latency M/NM selectivity in PFC than in PPC in 52 of 58 recording sessions from both monkeys (PFC : PPC, monkey Q: 151.2 ms : 199.1 ms, monkey Q: 157.3 ms : 171.7 ms, the difference in 46 sessions reached statistical significance P < 0.05, Wilcoxon test; **Figure S3D-G**). Together, the shorter latency M/NM selectivity in PFC compared with PPC is consistent with a preferential role for PFC in M/NM decisions.

**Figure 3.**
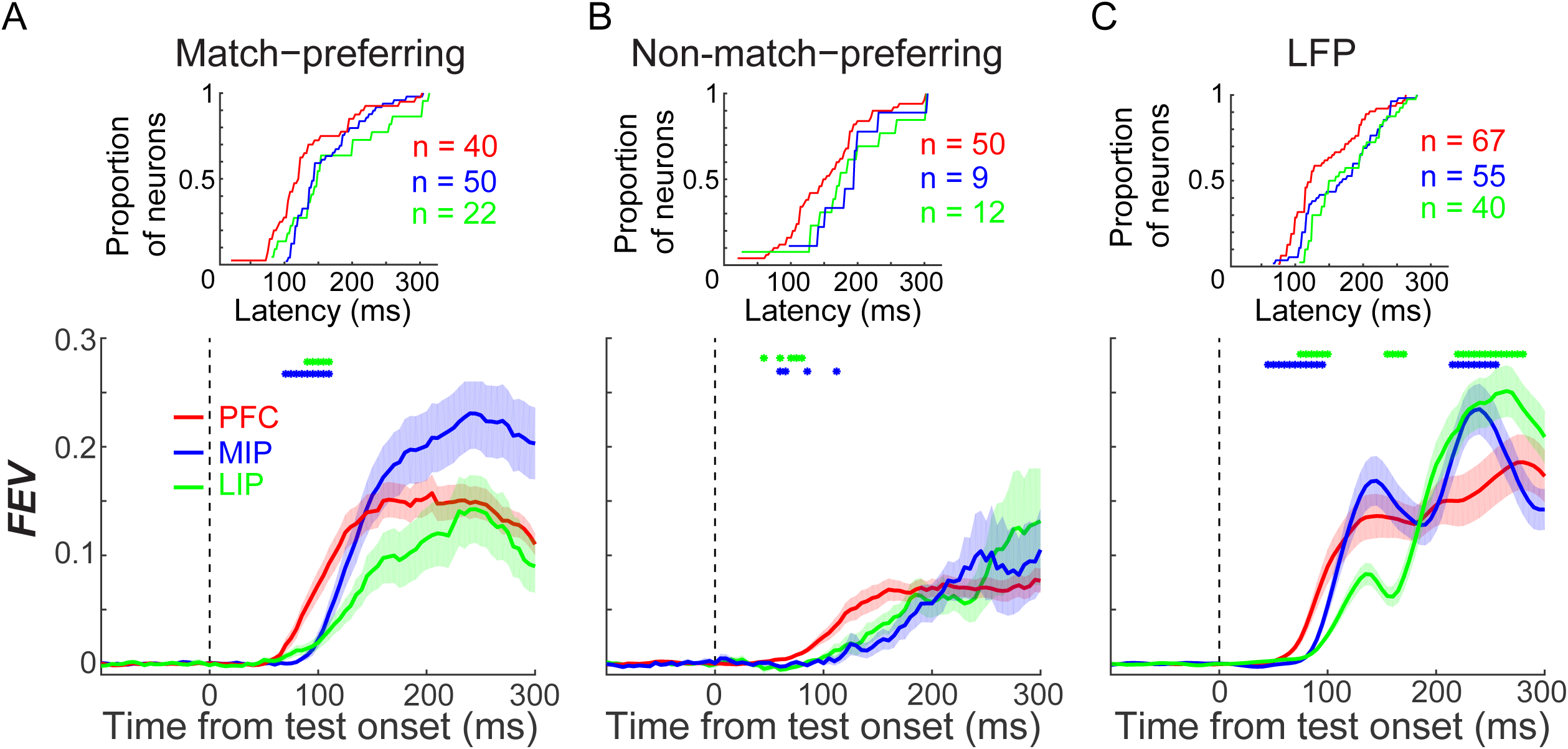
The comparison of M/NM selectivity between in PFC, LIP and MIP. **A-B**. The magnitude and time course of M/NM selectivity was determined using unbiased FEV. Different colors represent different cortical areas, and shaded area represents ±SEM. The blue dots denote the time points for which there were significant differences between PFC and MIP, while the green dots denote the time points for which there were significant differences between PFC and LIP (P < 0.05, Wilcoxon test). The upper insert figures show the cumulative distribution of the latency of M/NM selectivity. **C**. The M/NM selectivity of LFP amplitude in PFC, LIP and MIP, which is shown in the same format as (A). The LFP signal from all recording channels in each area are included.

Second, we tested whether the M/NM selectivity observed in each cortical area correlated with specific cognitive processes, such as the comparison of sample and test categories, or movement planning/initiation. For this analysis, we compared the activity of match-preferring neurons between match and non-match trials aligned to the monkeys’ lever release. In the DMC task, monkeys released the lever during both the first test period of match trials and the second test period of non-match trials. However, the decision about the match-status of the test stimulus occurred during the first test period of both types of trials, since the second test stimulus was only shown on non-match trails and was always a match (requiring a lever release). Match-preferring neurons in PFC showed greater activity preceding and up to the time of the lever release on match trials compared to non-match trials (200-0 ms prior to the hand movement, **Figure 4A**, p = 0.0018, paired t-test). In LIP, match-preferring neurons showed greater activity during match trials vs. non-match trials only prior to, but not coincident with, the hand movement (200-100 ms prior to the hand movement, **Figure 4B**, p = 0.0024; −100-0 ms, p = 0.21; paired t-test). The activity of match-preferring neurons in MIP during both match and non-match trials was very similar both before and during the hand movement (200-0 ms prior to the hand movement, **Figure 4C**, p = 0.99, paired t-test). These results suggest that match-preferring neurons in PFC and LIP are more involved in non-motor functions during M/NM decisions, such as the comparison of sample and test categories, while match-preferring neurons in MIP are primarily involved in motor functions such as planning and/or initiating hand/arm movements.

**Figure 4.**
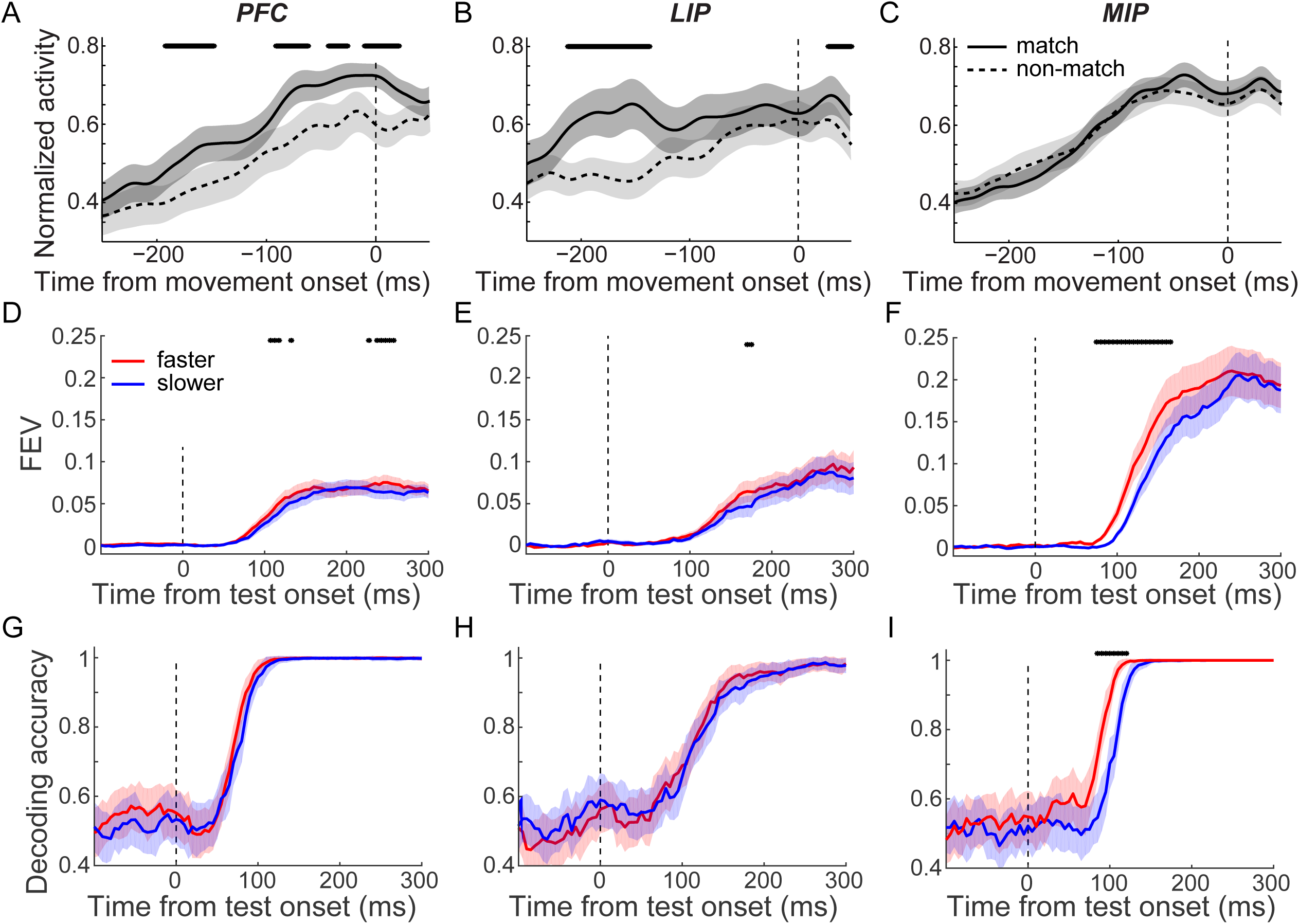
M/NM selectivity in PFC, LIP and MIP correlated with different cognitive processes during M/NM decision. **A-C**. The population activity of match-preferring neurons during match and non-match trials when activity was aligned to the start of hand movement. The black stars mark the time periods for which there were significant differences (p < 0.01, paired t test). **D-F**. The time course and magnitude of the M/NM selectivity in faster trials (red) and slower trials (blue) was evaluated using unbiased FEVs for PFC (D), LIP (E), and MIP (F). The shaded area represents ±SEM, and the black dots denote the time points for which there were significant (p < 0.01, paired t-test). **G-I**. The decoding performances for M/NM choices of an SVM classifier in faster (red) and slower trials (blue) for PFC (G), LIP (H), and MIP (I). The shaded area represents ±STD, and the black dots mark the time point when there were significant differences (p < 0.05, bootstrap).

Furthermore, we examined how neuronal M/NM selectivity covaried with the monkeys’ reaction time (RT) across the three cortical areas. To examine this, we separated match trials into two equal-sized RT sub-groups (fast and slow) for each neuron (fast: slow: LIP = 240.6 : 287.3 ms, MIP = 243.9 : 293.1 ms, PFC = 233.4 : 278.3 ms, see Materials and methods). We then compared M/NM selectivity for these sub-groups in each area. Both the unbiased FEV and the SVM analyses revealed significantly shorter-latency M/NM selectivity in MIP for faster vs. slower RT trials (**Figure 4F,I**, P_(FEV)_ = 0.0011, paired t-test; P_(SVM)_ < 0.02, bootstrap), but not in PFC or LIP (**Figure 4D,E,G,H**, PFC: P_(FEV)_ = 0.20, paired t-test; P_(SVM)_ > 0.3, bootstrap; LIP: P_(FEV)_ = 0.60, paired t-test; P_(SVM)_ > 0.40, bootstrap). Given the longer latency of M/NM selectivity in MIP and the preponderance of neurons preferring match conditions (which were accompanied by arm/hand movement), these results further suggest that M/NM selectivity in MIP is primarily associated with motor aspects of the decision.

### Integrative sample and test category representations in PFC and LIP

To solve the DMC task, the category membership of both sample and test stimuli need to be compared or integrated to form the M/NM decision. To gain insight into the basis for this integration, we examined how test-period activity in the three cortical areas encoded the previously-presented sample category and the currently-visible test category, and how sample and test category representation is related to the M/NM decision process. We first quantified the neuronal representation of sample and test categories in each area during the first test period. We focused on the time window preceding the mean RTs of both monkeys (0-250 ms after test onset), since sample and test category information must be integrated before the monkeys’ M/NM choice. Test period activity in LIP showed significantly stronger encoding of both sample and test categories than PFC and MIP (**Figure 5-figure supplement 1**, P_(LIP vs. PFC, sample)_ = 0.0083, P_(LIP vs. MIP, sample)_ = 0.00076, P_(LIP vs. PFC, test)_ = 0.0094, P_(LIP vs. MIP, test)_ = 8.3×10^−6^, Wilcoxon test)— consistent with our previous findings (Swaminathan and Freedman, 2012; Swaminathan et al., 2013)— suggesting that LIP is more directly involved in category computation than PFC. Meanwhile, test period activity in both PFC and LIP both showed a combined encoding of sample and test category information; but this was not observed in MIP (**Figure 5A-C**, PFC: FEV_(sam)_ = 0.0399, p = 6.7×10^−8^, FEV_(tes)_ = 0.0108, p = 1.7×10^−5^; LIP: FEV_(sam)_ = 0.1143, p = 3.5×10^−6^, FEV_(tes)_ = 0.0402, p =5.7×10^−4^; MIP: FEV_(sam)_ = 0.0230, p = 3.5×10^−5^, FEV_(tes)_ = 0.0026, p = 0.1146; paired t-test). This raises a question about the manner by which sample and test information is simultaneously encoded in both PFC and LIP—specifically whether sample and test information are encoded independently (e.g. additively), reflecting linear-like integration. Alternatively, sample and test information might be combined in a more nonlinear fashion in either or both areas, reflecting local computation.

**Figure 5.**
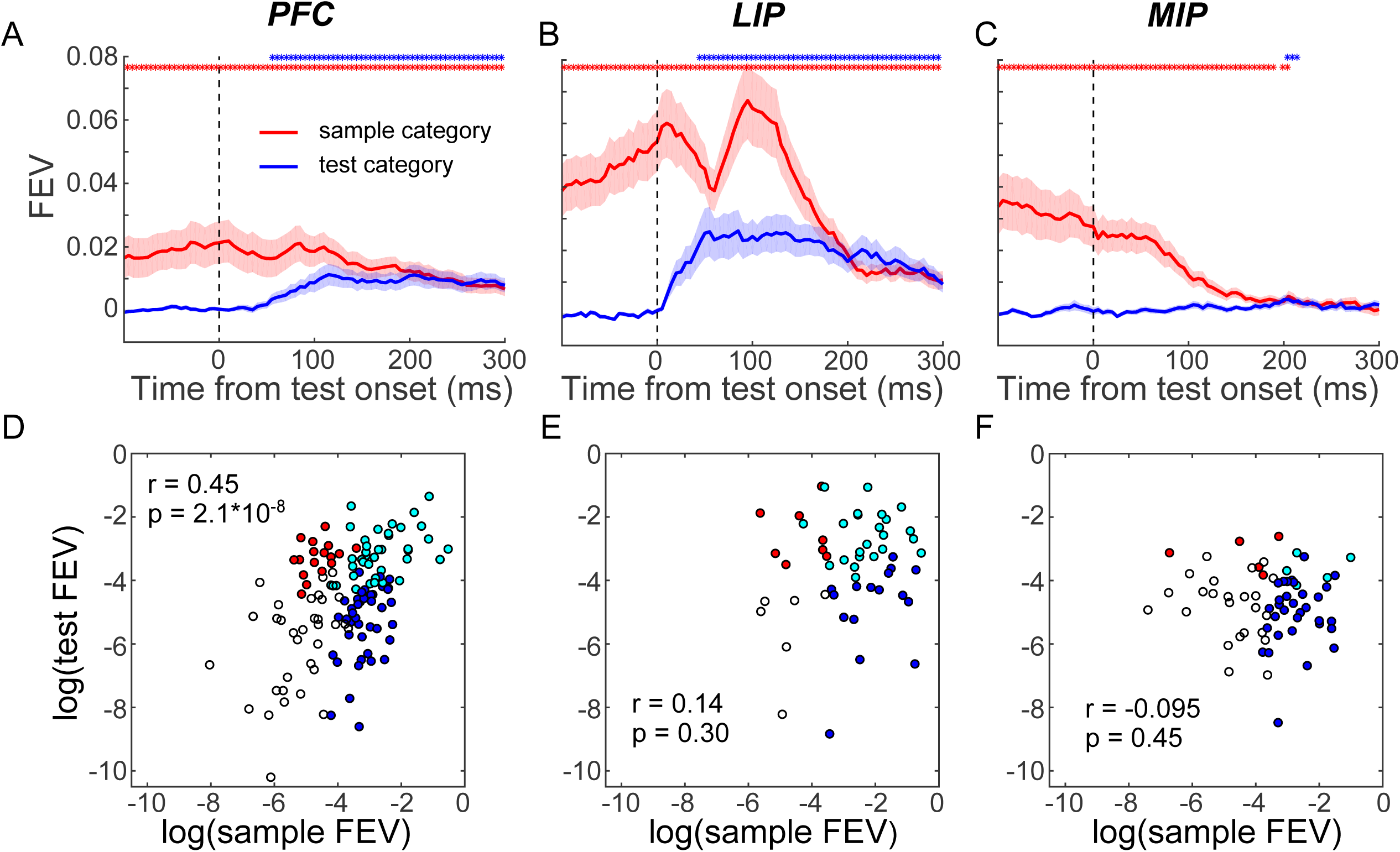
Sample and test category representation in PFC, LIP and MIP. **A-C**. The selectivity of sample category (red) and test category (blue) were evaluated using the unbiased FEV for all neurons in PFC (A), LIP (B) and MIP (C). The shaded area represents ±SEM. The red and blue dots represent the time points for which the sample and test category selectivity are significantly greater than chance level (P < 0.01, paired t-test), respectively. **D-F**. The correlations between sample category and test category selectivity (using FEV) during the test period for all the neurons in PFC (D), LIP (E) and MIP (F). Each symbol represents a single neuron. Cyan dots denote neurons that showed significantly mixed sample-test-category selectivity. The blue and red dots denote the neurons that showed only significant sample category or test category selectivity respectively, and the black circles denote the neurons that did not show significant category selectivity (one-way ANOVA test, p < 0.01).

To better understand how PFC and LIP integrate sample and test category information, we examined how each area simultaneously encoded the sample and test categories during the test period. First, we asked whether there was a correlation between the strength of neurons’ sample and test category encoding. A positive correlation between sample and test category representations would suggest that a special pool of neurons is preferentially involved in encoding both the remembered sample and currently visible test categories, indicative of sample-test category integration at the single neuron level. In contrast, a negative correlation or zero correlation would indicate that there is less overlap of the neurons’ sample and test category encoding. We calculated the unbiased FEV of sample and test category encoding for each neuron’s test-period activity and found PFC and LIP neurons that showed both sample and test category selectivity. However, the correlation between sample and test category representations at the population level differed between PFC and LIP. In PFC, there was a significant positive correlation between sample and test category selectivity shortly after test onset (100-200 ms,**Figure 5D**, r = 0.3719, P < 0.0001, t-test), while in LIP, they were not correlated (**Figure 5E**, r = 0.1152, p = 0.4115, t-test). The positive correlation in PFC was also evident by using the category tuning index (rCTI) to quantify category selectivity (r = 0.3323, p = 0.001) (see Materials and methods). Furthermore, neuronal sample category selectivity prior to test onset in PFC but not LIP (-50-50 ms relative to test onset) was correlated with M/NM selectivity during the test period (150-250 ms after test onset) (PFC: r = 0.2808, p = 0.0006; LIP: r = −0.0420, p = 0.7654), suggesting that PFC neurons with greater sample category encoding before test onset were more likely to be involved in the M/NM computation. These results suggest that single neurons in PFC, but not in LIP, integrate sample and test category information. MIP is unlikely to be involved in such an integration process, as very few neurons showed significant encoding of test category (**Figure 5F**).

Next, we examined whether sample and test category selectivity was integrated in a linear or nonlinear manner in PFC and LIP. In the DMC task, there were four possible combinations of sample and test categories (i.e. S_1_T_1_, where both sample and test stimuli are category 1, S_1_T_2_, S_2_T_1_, and S_2_T_2_). Examining the correlation of test category selectivity between the two sample category conditions (S_1_T_1_ vs. S_1_T_2_ and S_2_T_1_ vs. S_2_T_2_) provides insight into the way in which sample and test categories are integrated. We attempt to differentiate between two possible outcomes, with each suggestive of a particular kind of representation: (1) test category selectivity that added linearly to the existing sample category selectivity, in which the test category selectivity in the two sample category conditions would be similar in sign and magnitude (positive correlation) (2) test category selectivity that combined nonlinearly with sample selectivity, in which the test category selectivity for the two sample category conditions would be different (negative or no correlation). In particular, a negative correlation would result if the neuron shows an opposite test category preference between the two sample category conditions (i.e., M/NM selectivity). We evaluated these possibilities in PFC and LIP using two approaches. The first was an ROC analysis which quantified, for each neuron, test category selectivity in each of the two sample category conditions, and then calculated their correlation across the population in each area. This revealed a positive correlation in LIP shortly following test onset **(Figure 6B** maximum r = 0.7052, p < 0.001, t-test), but not in PFC **(Figure 6A** maximum r = 0.2472, p = 0.0027). Second, we trained a decoder (SVM) to classify test category using the trials in one sample category condition (e.g. S_1_T_1_ and S_1_T_2_), and tested the decoder using trials from the other sample category condition (e.g. S_2_T_1_ and S_2_T_2_). Decoder performance is expected to be above chance if the test category selectivity was similar in each of the two sample category conditions, or below chance if the test category selectivity differed based on which sample stimulus has been shown on that trial. As shown in **Figure 6C-D**, decoder accuracy is significantly greater than chance shortly after test onset in LIP (maximum accuracy = 0.7738, p < 0.01, bootstrap), but does not exceed chance in PFC (maximum accuracy = 0.5381, p > 0.50, bootstrap). Together, these results suggest that the combination of sample and test category information during the test period is more consistent with linear integration in LIP but not PFC.

**Figure 6.**
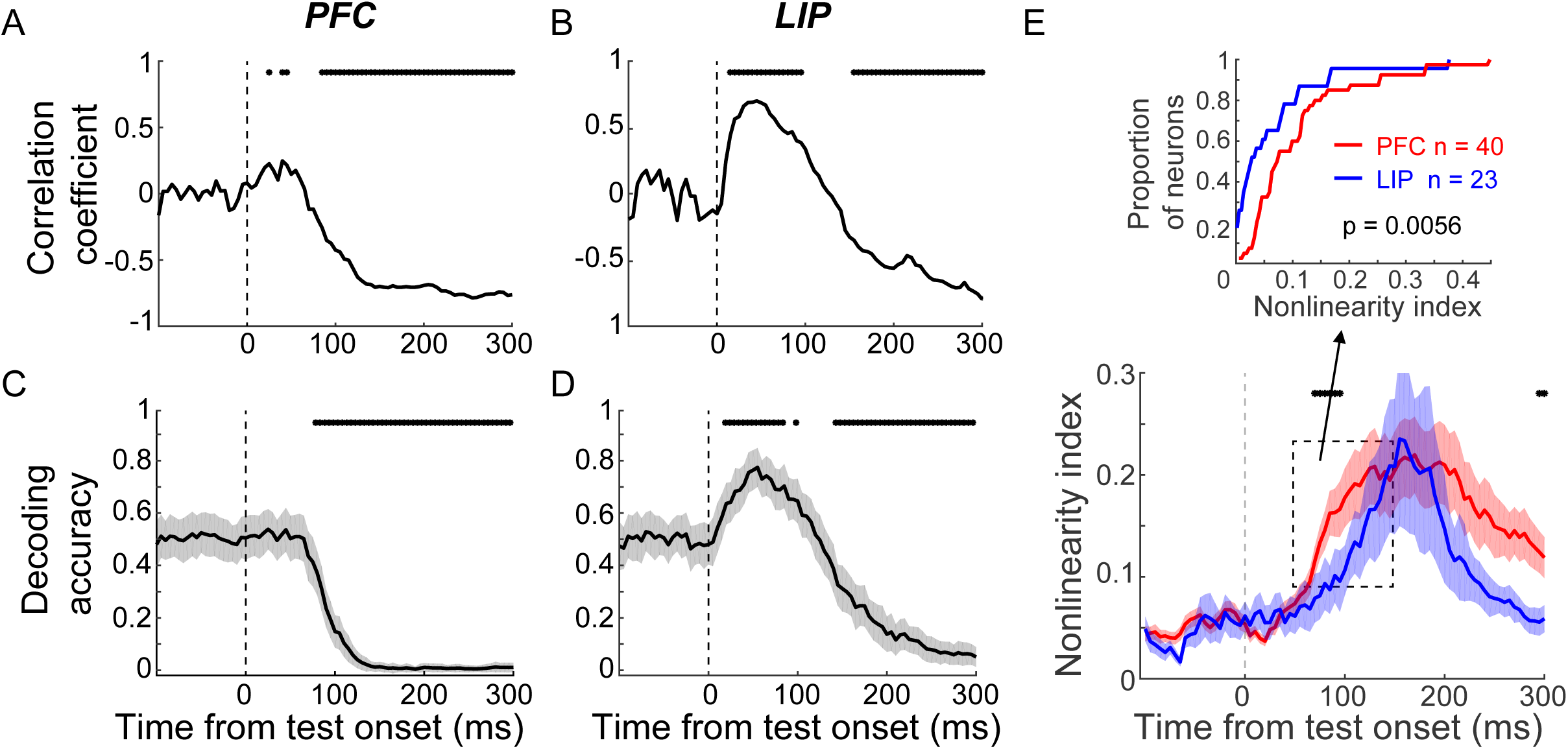
The mixed category selectivity was more nonlinear in PFC than in LIP. **A-B**. The correlation coefficient between the test category selectivity (using ROC value) of two sample category conditions is shown for both PFC (A) and LIP (B). The black dots mark the time points for which the correlation was statistically significant (p < 0.01, t-test). **C-D**. The decoding performance of a test category classifier using neuronal activity in PFC (C) and LIP (D). The SVM classifier was trained by using activity from one sample category condition (e.g. S1T1 vs. S1T2), and tested with activity from the other sample category conditions (e.g. S2T1 vs. S2T2). The shaded area represents ±STD, and the black stars mark the time points for which the decoder performance is significantly different from chance level (bootstrap, p < 0.05). **E.** The nonlinearity index of mixed category selective neurons in both PFC (red) and LIP (blue). The shaded area represents ±SEM, and the black stars denote the time points for which there is a significant difference between LIP and PFC (Wilcoxon test, p < 0.05). The upper panel shows the cumulative distribution of nonlinearity index shortly after test onset (50-150 ms after test onset) for both PFC and LIP neurons.

We quantified the degree to which the integrated sample-test category encoding during the test period was linear or nonlinear by calculating a nonlinearity index for LIP and PFC neurons which showed both sample and test category selectivity in both LIP and PFC. The nonlinearity index was defined as the absolute difference between the test category selectivity in two sample category conditions (quantified by the absolute mean activity difference between S_1_T_1_ and S_1_T_2_, or S_2_T_1_ and S_2_T_2_), which was then normalized by the sum of mean activity in the four sample-test-category combinations (see Materials and methods). The value of the nonlinearity index, which is not expected to be affected by linearly combined category selectivity or M/NM selectivity, can range from 0 to 1. Values near 0 indicate linear-like selectivity of the two factors while increasing values indicate nonlinear combination of sample and test category selectivity. Because the neuronal activity shortly before monkeys’ M/NM choice mainly correlated with monkeys’ M/NM choice (**Figure 6**), we focus on the time window shortly after test onset (50-150 ms). As shown in **Figure 6E**, the nonlinearity index is significantly greater in PFC than LIP during the early test period (p = 0.0056, Wilcoxon test). This suggests greater nonlinearity in the combination of sample and test category representation in PFC compared to LIP, independent of the strength of M/NM selectivity in these two areas. These results further confirm that integrated sample-test category representations are more nonlinear in PFC than LIP. The more linearly integrated sample and test category representations in LIP suggest that it is better suited for independently encoding the remembered sample stimulus and currently visible test stimulus, perhaps facilitating readout of these variables by downstream cortical areas. Considering the observation of shorter-latency M/NM selectivity in PFC than LIP and MIP (**Figure 3**), the positively correlated and more nonlinear integration of sample-test category selectivity in PFC is consistent with it combining remembered sample and visible test category information in order to facilitate M/NM decisions.

### Nonlinear PFC encoding was preferentially engaged in M/NM decisions

The results presented so far suggest that PFC is more involved in integrating sample and test category information to form M/NM decisions, compared to LIP and MIP. We tested this idea more directly by assessing the relationship between PFC neurons’ activity and monkeys’ M/NM decisions as a function of the linearity or nonlinearity of their selectivity for sample and test categories. We performed a two-way ANOVA on test-period activity with sample and test categories as factors. This allowed us to identify two populations of PFC neurons: (i) linearly integrating neurons (LIN), which exhibited main effects of both sample and test categories (p < 0.01) but a non-significant interaction term; and (ii) nonlinearly integrating neurons (NIN), which exhibited main effects of both sample and test categories (p < 0.01), as well as a significant interaction term (p < 0.01). We also identified pure-selective neurons (PN), which showed significant effect of only sample category, test category or interaction. Note that the M/NM selective PNs are different from the NINs as they showed neither sample nor test category selectivity. **Figure 7A-C** shows test-period activity of three NINs, each of which encoded both sample and test categories, and preferentially responded to one of the four sample-test category combinations. In order to test whether the NINs were more involved in mediating DMC task performance than the other groups of PFC neurons, we compared the strength with which three key task variables (sample category, test category, M/NM) were encoded among the NINs, LINs, and PNs using the unbiased FEV **Figure 7D-E**). Interestingly, NINs showed significantly stronger sample category encoding than either the LINs or sample-category selective PNs in PFC (**Figure 7D**, P_(NIN vs. LIN)_ = 0.0036; p_(NIN vs. PN)_ = 4.6×10^−8^; Wilcoxon test), while the strength of sample-category selectivity did not differ between LINs and sample-category selective PNs (P = 0.0917, Wilcoxon test). Furthermore, NINs showed significantly greater and shorter latency M/NM selectivity than the M/NM selective PNs in PFC (**Figure 7E-F**, p(latency) = 8.7×10^−4^, p(magnitude) = 5.6×10^−4^, Wilcoxon test). Since LINs did not show significant M/NM selectivity based on our criteria, we did not include them for this analysis. To ensure that this difference between NINs and other PFC neurons was not due to differences in firing rates among the different groups, we compared the mean test-period activity among these groups of neurons and found non-significant differences (NIN: 12.8sp/s ; LIN: 11.0sp/s, p_(NIN vs. LIN)_ = 0.9670; category selective PN: 12.2sp/s, p_(NIN vs. PN)_ = 0.7730; M/NM selective PN: 11.9sp/s, p_(NIN vs. PN)_ = 0.7529, Wilcoxon test). However, NINs did not show significantly greater test category selectivity than LINs and test-category selective PNs (p_(NIN vs. LIN)_ = 0.4642, p_(NIN vs. PN)_ = 0.2235). This might be because PFC showed weaker test category selectivity compared to LIP, and may therefore be less involved in rapidly encoding the currently visible test category compared to LIP.

**Figure 7.**
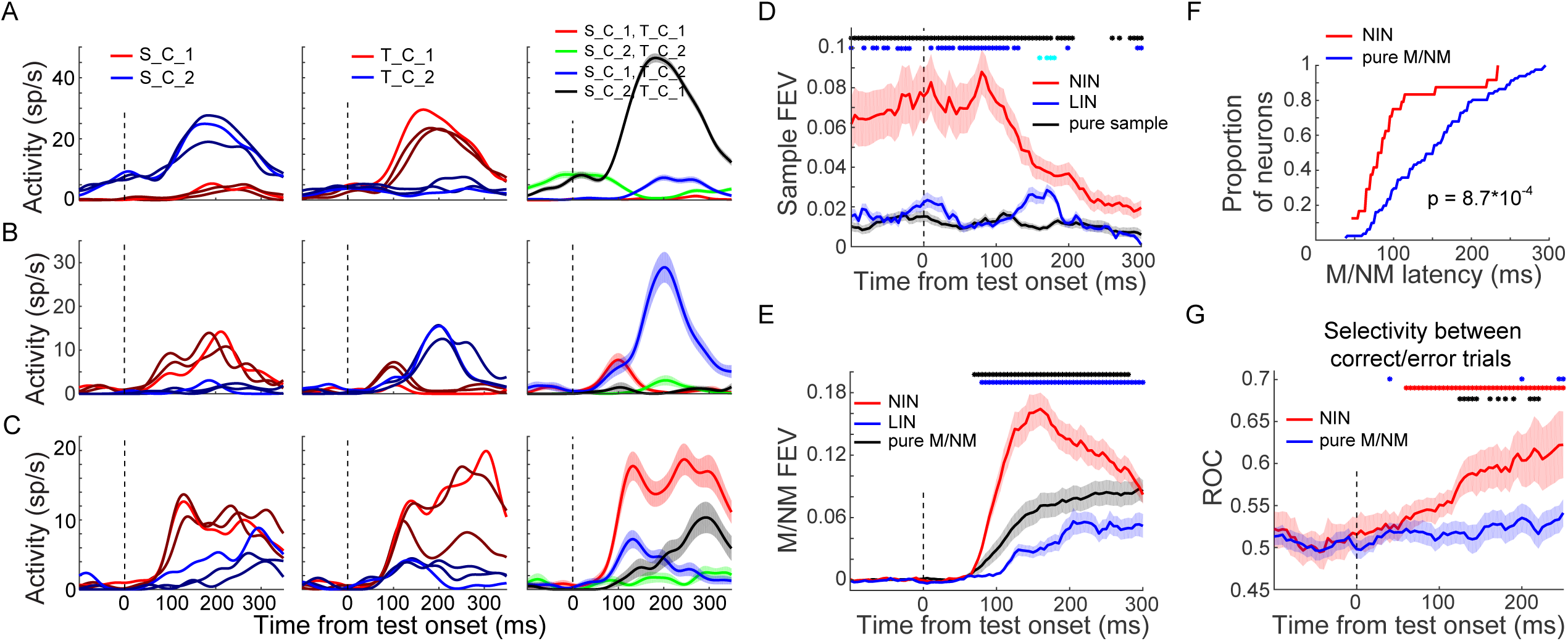
The nonlinearly integrating neurons (NIN) in PFC were more engaged in the DMC task. **A-C**. The activity of three NINs. The first and second columns show the condition-averaged activity sorted by sample direction and test direction, respectively. The lighter colors correspond to directions in the center of each category and dark colors correspond to all other directions. The third column shows the mean activity of each sample-test-category combination. The shaded area denotes ±SEM. S: sample, T: test, C: category. **D**. The sample category selectivity of the NIN, linearly integrating neurons (LIN) and pure-sample-category-selective neurons in PFC were compared using FEV. The shaded area denotes ±SEM. The blue and black dots denote the time point for which the NINs were significantly different from LINs and pure-sample-category-selective neurons, respectively; while the cyan dots denote the time point for which there was significant difference between LIN and pure-sample-category-selective neurons (p < 0.05, Wilcoxon test). **E**. The M/NM selectivity of the NINs, linear neurons and pure-M/NM-selective neurons in PFC were compared using FEV. The color dots denote the statistical significance in the same format as in d. **F.** The cumulative distribution of the latency of M/NM selectivity for NINs and pure-M/NM-selective neurons. **G**. The activity change of incorrect-match trials relative to correct-match trials were evaluated using ROC for both NINs and pure-M/NM-selective neurons. The shaded area denotes ±SEM. The red and blue dots denote the time points for which the activity changes of NINs and pure-M/NM-selective neurons were statistically significantly (p < 0.05, paired t-test), respectively; while the black dots denote the time points for which there were significant difference between NINs and pure-M/NM-selective neurons (p < 0.05, Wilcoxon test).

To test whether the activity of NINs was more correlated with the monkeys’ M/NM decisions compared to M/NM-selective PNs in PFC, we compared neuronal activity on correct and incorrect match trials (in which monkeys should have released the lever in response to the first test stimulus; monkeys made very few errors on non-match trials) with an ROC analysis (see Materials and methods). ROC values greater than 0.5, indicate that a neuron’s M/NM selectivity covaries with the monkey’s trial-by-trial M/NM choices, while values near or lower than 0.5 indicate no correlation or anti-correlation, respectively. As shown in **Figure 7G**, significantly elevated ROC values indicate that the activity of NINs (but not M/NM-selective PNs) reflected the monkeys’ trial-by-trial M/NM decisions (NIN: ROC = 0.604, P = 0.0019; PN: ROC = 0.529, P = 0.4325, paired t-test). Furthermore, the difference in NIN activity between correct and incorrect match trials (measured by ROC) was greater than that of M/NM-selective PNs neurons (P = 0.0178, Wilcoxon test), suggesting that activity of NINs was more closely related to the monkey’s trial by trial M/NM decisions. Together, our results suggest that PFC neurons which showed nonlinear integration of sample and test category information play a preferential role in the formation of M/NM decisions.

### PFC NINs are crucial for solving the M/NM computation in silico

RNN models trained on complex behavioral tasks have been recently shown promise as an approach to understand neural computations (Engel et al., 2015; Masse et al., 2019; Song et al., 2016) — particularly for behavioral tasks which require integrating or comparing events across time. We therefore trained RNNs to perform the DMC task in order to further test the idea that nonlinear integrative encoding in PFC plays a critical role in sequential decisions. Recent studies from a number of groups, including our own, have employed RNN models with a single pool of recurrent units in the hidden layer. However, this poses a challenge for relating modeling work to neuroscience questions involving multiple interconnected brain areas. To address this challenge, we developed a novel approach to a multi-module RNN, in which the modularity and rules governing the connections between modules were inspired by neurobiological principles. These RNNs are composed of two hierarchically organized modules, with the module closer to sensory input intended to correspond to LIP, and the module closer to the motor output corresponding to PFC. The modularity and hierarchy were imposed through a set of initial constraints on the recurrent weight matrices, as well as the input and output weight matrices (**Figure 8A-B**, see Materials and methods). These constraints were designed to emulate the architectonic principles that govern intracortical projections. Both modules were assigned 50 percent of units in the network, with matched proportions of excitatory/inhibitory neurons (80% excitatory to 20% inhibitory). The LIP module receives the motion direction input, while the PFC module projects to two response units which simulate holding and releasing touch-bar respectively. Aside from these biologically-derived architectural features, the modules’ functional roles were not explicitly constrained (e.g. the LIP module and PFC module were not forced to encode category information and decision information, respectively). We independently trained 50 networks with randomly initialized weights and identical hyperparameters to perform the DMC task, using methods previously described (Masse et al., 2019).

**Figure 8.**
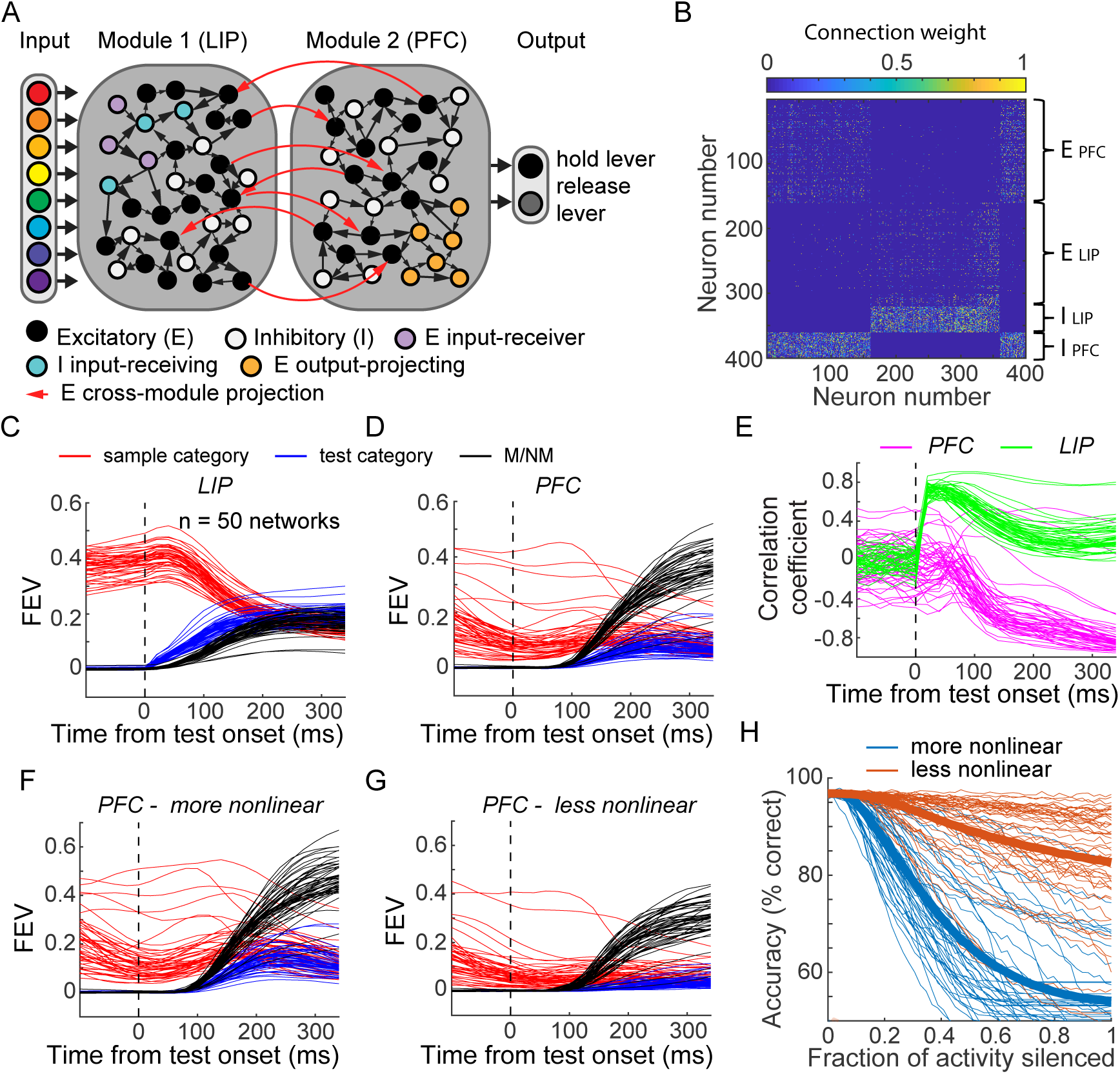
PFC nonlinear integration is crucial for mediating the M/NM decisions in RNNs. **A**. Model schematic of the two-module ‘frontoparietal’ RNNs. Each RNN is consist of 24 motion direction turned input units, 400 hidden units and two response units. The hidden layer of each RNN consists of two modules simulating LIP and PFC respectively, with half of the units designated to each module (and E/I proportion maintained). A subset of LIP units, which have recurrent connections only within module, receive the motion tuned inputs; while another LIP excitatory units sparsely project to all units in PFC module, except for the excitatory subset that project directly to the output units. The subset of PFC excitatory units that do not project to the output also send sparse feedback projections to LIP module. Both the excitatory and inhibitory units in each module are recurrently connected within each module. **B**. Example recurrent connectivity matrix of an example two-module RNN. Inhibition is strictly local to each module, as is emphasized by the block-diagonal structure in the bottom fifth of rows. Excitatory projections between modules are sparse, while excitatory projections within modules are denser. **C**. The averaged sample category selectivity, test category selectivity and M/NM selectivity of units in the LIP modules of the 50 RNNs were quantified using FEV. Each line denotes the result from one RNN. **D**. The sample category selectivity, test category selectivity and M/NM selectivity of PFC modules. **E**. The correlation coefficient between the test category selectivity of two sample category conditions is shown separately for both PFC (pink) and LIP (green) modules. **F-G**. The task variable encodings (sample, test and M/NM) of the more-nonlinear (F) and less-nonlinear (G) groups of units in the PFC module are shown separately. **H**. The behavior performances of the RNNs after gradually inactivating the more-nonlinear and less-nonlinear groups of units in the PFC module. The thick lines denote the averaged performances of all 50 RNNs.

After training, all 50 networks converged to perform the DMC task with high accuracy (95.3% ± 0.78%). Both LIP and PFC modules’ units encoded all three key task variables during the test period in a similar manner to the neurophysiological data (**Figure 8C-D**). LIP module units showed significantly greater encoding of both sample and test categories (sample: p = 2.2e-13, tstats = 9.98; test: p = 1.8e-37, tstats = 37.0133; df = 49, paired t test), while PFC module units showed significantly greater N/NM encoding (p = 1.9e-27, df = 49, tstats = 22.45, paired t-test) Individual units in both LIP and PFC modules encoded both sample and test category (**Figure 8-figure supplement 1**). As in the real LIP and PFC data, sample and test category encoding in the RNNs during the test period was more strongly correlated in the PFC than LIP module (r_PFC_ = 0.41± 0.16, r_LIP_ = 0.057± 0.097, p = 1.7e-21, df = 49, tstats = 16.37, paired t-test). Furthermore, sample and test category encoding were more likely to be nonlinearly integrated in the PFC than the LIP module, shown by examining the correlation of test category selectivity between the two sample category conditions (**Figure 8 E**). Furthermore, the more-nonlinear units in PFC module showed greater encoding of all the key task variables compared to other task related units (**Figure 8F-G**, sample: p = 6.0071e-23, tstats = 17.7183; test: p = 3.8848e-30, tstats = 25.7542; M/NM: p = 4.3940e-25, tstats = 19.8589, df =49, paired t-test). These results suggest that neural activity and information encoding in the two-module RNNs closely resembled the neurophysiological data, and lends support to the idea that nonlinear encoding of task-related variables in neural networks close to the output of the decision process (i.e. both PFC and the higher order RNN module) are critical for mediating sequential decisions.

To directly test whether nonlinear units in the PFC module of the RNNs play a causal role in M/NM decisions, we performed an inactivation study *in silico*. We selectively silenced different groups of units in the PFC module during the test period, and then tested the impact of that inactivation on behavioral performance of the network. To do this, we first computed the nonlinearity index for all task-related units in the PFC module for each RNN. After leaving out the units directly projecting to the response units, we then silenced the top 50% of units (nonlinear group) and the bottom 50% of units, ranked by nonlinearity index. This procedure allows a direct test of the causal involvement of different PFC units based on their degree of nonlinear sample-test encoding. Specifically, we performed precise, graded inactivation -- smoothly titrating the amount of silencing applied to each unit from complete (100% reduction in activity) to minimal (0% reduction in activity). At every level of inactivation, the nonlinear group resulted in a more severe behavioral effect than the other group, even though they contained the same number of neurons and were inactivated to the same extent (**Figure 8 H**). This implicates a specific ablation of the M/NM computation, rather than a gross disruption of the network’s wiring to decision readouts, in the behavioral deficit that results from inactivating the nonlinear group of PFC units. Additionally, this difference was not due to differences in activity level of the two groups of units, as the mean activity was similar between the two groups (p = 0.7459, tstats = 0.3258, df = 49, paired t test) during the test period. Together, the two-module RNN simulations and *in silico* inactivation experiments reinforce the plausibility of our central neurophysiological findings of the importance of nonlinear encoding of task-relevant information for mediating sequential decisions during the DMC task.

## Discussion

This study examined PFC, LIP and MIP activity in an effort to illuminate their roles in the sensory, cognitive, and motor aspects of decision making during a DMC task which required monkeys to indicate whether sequentially presented sample and test stimuli were from the same (matching) or different (non-matching) categories. In particular, we sought to understand the mechanisms by which the remembered sample stimulus and currently visible test stimulus are integrated to reach a M/NM decision. We found that although test-period activity in PFC, LIP and MIP all correlated with the monkeys’ M/NM decisions, these areas play distinct roles in the decision process. Our results suggest that PFC functions as a candidate source of M/NM decision signals, which are likely generated by nonlinearly integrating inputs from upstream areas into encoding of decision-related task variables. LIP appears to be primarily involved in the rapid analysis of stimulus category during stimulus presentation, as well as encoding previously viewed stimuli in short term memory. In contrast, MIP activity primarily reflects the monkeys’ planned and ongoing motor responses (i.e. the hand movements used to report their decisions). These results elucidate differential and distributed roles of frontal-parietal network nodes during sequential decisions, and suggest that nonlinear-integration of task-related variables in PFC is a key mechanism for sequential M/NM computations.

In order to correctly decide whether a test stimulus matches the category of a previous sample, the brain must accomplish several tasks. First, it must correctly categorize the sample stimulus, and maintain a representation of sample category throughout the delay and into the test period. The brain must then correctly identify the test stimulus category, compare the test category to the remembered sample category, and select the appropriate behavioral response to report the M/NM decision. Previous work from many groups, including our own, has identified encoding in PPC and PFC related to these various task components. For example, both PPC and PFC can encode the identity and task-relevance of sensory stimuli, and maintain such information in neuronal activity during short-term memory delays (Cromer et al., 2010; Freedman and Assad, 2006; Freedman et al., 2001; Sarma et al., 2016; Swaminathan and Freedman, 2012). Furthermore, PFC and PPC activity correlates with a wide-range of decision processes, including perceptual, categorical, and M/NM decisions (Cromer et al., 2011; Crowe et al., 2013; de Lafuente et al., 2015; Ding and Gold, 2012; Freedman and Assad, 2006; Kiani and Shadlen, 2009; Mante et al., 2013; Platt and Glimcher, 1999; Qi et al., 2012; Sugrue et al., 2004).

By directly comparing PFC, LIP, and MIP activity in the same animals performing a DMC task, our results give new insight into the relative functions of each area in both categorical and M/NM decisions, and into the format of neuronal encoding which may underlie computation of M/NM decision signals. We show that test-period activity in LIP showed the strongest categorical encoding of both the remembered sample and the currently visible test stimulus. This supports the idea that LIP is involved in categorical representation of both visible and remembered stimuli during the DMC task. In contrast, LIP appears less involved in transforming categorical encoding into M/NM decisions. This is supported by the longer-latency M/NM selectivity in LIP compared with PFC. On the other hand, although test-period activity in MIP reflects the remembered sample category, we did not detect an obvious encoding of the currently visible test category in MIP. Instead, MIP activity during the test period was dominated by motor-related encoding arising with a longer latency compared to M/NM selectivity in PFC, consistent with previous work (Cui and Andersen, 2007; Swaminathan et al., 2013). These results indicate that MIP is unlikely to be directly involved in the comparison of sample and test categories, but instead primarily associated with motor aspects of the tasks, such as planning the movements used to report the monkeys’ decisions.

Three lines of evidence suggest that PFC, rather than LIP and MIP, is a leading candidate for guiding M/NM decisions during sequential decision tasks such as the DMC. First, M/NM selectivity of both spiking and LFP signals arose with a shorter latency in PFC than in both PPC areas, suggesting a flow of M/NM encoding from PFC to PPC. Second, PFC neurons showed a relatively balanced preference for both ‘match’ and ‘non-match’, while LIP and MIP were biased toward preferring ‘match’ conditions, which were accompanied by hand movements. Balanced M/NM representation in PFC suggests it is more likely to reflect the abstract M/NM decision rather than preparatory motor activity. Furthermore, those PFC neurons responding more strongly to non-matching test stimuli may be involved in PFC’s established role in response inhibition (Aron et al., 2014; Kramer et al., 2013; Schall and Godlove, 2012) (i.e. withholding a motor response on non-match trials). Third, the sample and test categories are combined in a nonlinear manner in PFC, but not in LIP or MIP. This nonlinear encoding could endow the PFC network with a high dimensional representation to integrate the remembered sample category and currently visible test category into a more task-related representation of the sample-test category combinations, which could then be mapped onto M/NM decisions. Together, these results are suggestive of a distinct M/NM decision process in PFC intervening between stimulus evaluation (motion categorization) in areas related to perceptual/decision processing such as LIP (Zhou and Freedman, 2019), and motor planning in motor-related areas such as MIP. This is consistent with previous reports of “abstract” rule encoding (i.e. independent of stimulus features or motor responses) in PFC and PPC (Stoet and Snyder, 2004; Wallis et al., 2001). Meanwhile, previous studies using a delayed match to sample task with visual motion stimuli observed comparison related activity in both PFC and medial temporal cortex (MT), but found that such activity was decision-correlated only in PFC (Lui and Pasternak, 2011; Zaksas and Pasternak, 2006). This is consistent with our finding that PFC is likely to be a source for mediating the abstract sequential decisions. Related findings extend beyond the visual sensory modality as well. For example, one line of work trained monkeys to compare sequentially presented vibrotactile stimuli, and found that neuronal activity in ventral premotor cortex reflected the process of comparing the sample and test stimuli (Romo et al., 2004; Rossi-Pool et al., 2016). Together, these results suggest that PFC may function as an abstract decision stage that integrates sequences of stimuli or events to determine their behavioral relevance, with nonlinear integration across task stimuli or events playing an important role in that process.

The frontal-parietal network has been implicated in mediating perceptual decision-making tasks which likely share cognitive demands and underlying mechanisms with the DMC task (Freedman and Assad, 2011, 2016). It was proposed that perceptual decision tasks are mediated by an “intentional framework”, in which decisions play out as a competition between representations of potential action plans (Gold and Shadlen, 2000, 2007). As we argued previously, it is unclear how an action-based framework can be easily extended to account for categorical decisions in the DMC task (Freedman and Assad, 2006, 2011), since the two categories are not rigidly linked with distinct actions. In contrast, M/NM decisions in the DMC task do map directly onto distinct motor responses (i.e. match = release lever; non-match = hold lever), raising the possibility that an intentional framework could potentially account for mechanisms underlying M/NM decisions. However, we interpret our results as favoring a more abstract (and less motor) framework. This is primarily because activity in MIP, a motor-related area showing activity closely correlated with the monkeys’ decision-related arm movements, appeared to be post-decisional and did not reliably encode key sensory and task-related variables needed to generate the M/NM decision. On the other hand, PFC activity was more consistent with playing a direct role in the M/NM decision process. However, PFC M/NM selectivity was less closely correlated with the RTs of monkeys’ motor responses compared to MIP. This suggests that the M/NM decision is unlikely to be achieved directly through sensorimotor transformation within the motor network, but might be generated through a distinct abstract M/NM decision stage. This could be tested by training monkeys to report their decisions using different effectors (i.e. saccade vs. manual response). The prediction is that M/NM selectivity in PFC would be closely correlated with monkeys’ M/NM decisions across effectors, whereas M/NM selectivity in the motor network would be effector specific.

Cortical neurons have been shown to encode mixed representations of multiple task variables during cognitively demanding tasks (Johnston et al., 2020; Parthasarathy et al., 2017; Rigotti et al., 2013; Zhang et al., 2017), and nonlinear mixed selectivity (NMS) in PFC has been particularly emphasized as an important mechanism for cognitive computations. Specifically, NMS can potentially facilitate a linear readout of task variables, and the strength of NMS is correlated with the subjects’ behavior (Fusi et al., 2016; Ramirez-Cardenas and Viswanathan, 2016; Rigotti et al., 2013). This is potentially related to the results of the current study. In LIP, sample and test categories were combined more linearly, and with selectivity for each variable randomly distributed across neurons. In contrast to LIP, sample and test category selectivity was more nonlinearly combined in PFC, and the strength of sample and test category encoding was positively correlated. In particular, the PFC neurons showing nonlinearly combined encoding of sample and test categories appeared to be most closely involved in the decision process. This is consistent with the idea that mixed selectivity in PFC and LIP may relate to the core functions of each area during the DMC task, and highlight NMS in PFC as playing a key role in cognitive computations for solving cognitive tasks. The two-way ANOVA approach which identifies linearly and nonlinearly integrated sample and test category representations used here is conceptually similar to the approaches used in the original NMS study. However, the structure of the task used in the current study prohibits us from performing a similar analysis as in the earlier NMS studies due to the smaller number of distinct conditions tested in our task (Fusi et al., 2016; Ramirez-Cardenas and Viswanathan, 2016; Rigotti et al., 2013).

Artificial neural networks have recently gained prominence as a model system for investigating neural mechanisms of cognition, together with neurophysiological experiments. Their advantages are largely complementary to those of neuronal experiments; in particular, they facilitate analyses of substrates that are difficult to measure and perturb in vivo. A recent study from our lab, using a machine-learning based training procedure to train RNNs, characterized the different roles of short-term synaptic plasticity and persistent spiking activity in maintaining and manipulating information within short-term memory (Masse et al., 2019). In the current study, we trained RNNs with a hierarchy of distinct modules, meant to mimic hierarchically interconnected cortical areas. Without explicitly specifying the functional roles of the modules before or during training, we enforced these modules from the beginning of training through a set of weight constraints which were designed based on the neurobiological principles of corticocortical connections. Based on the striking similarities between the way that units in these RNNs and LIP/PFC encode task variables, multi-module RNNs hold promise for modelling computations distributed across multiple brain areas, as well as for testing different areas’ relative roles during cognitive functions. Despite the technical difficulty of training biologically-inspired RNNs with multiple modules in past work, several recent efforts have begun to explore these models’ unique benefits to studies of cognitive functions. Our approach is most similar to that of Kleinman and colleagues (Kleinmana M. et al., 2019), who trained multi-area RNNs with separate E/I populations to perform a perceptual decision-making task. However, our approach differs from that study in three ways: 1) we incorporate more biological bottlenecks for information flow both within and between modules (units that directly receive bottom-up projections cannot themselves be a direct feedforward conduit out of their own module, and input for each module targets both excitatory and inhibitory units); 2) our networks are endowed with short-term synaptic plasticity, which affects the circuit mechanisms that emerge in the service of solving tasks; and 3) we evaluate our network in a different task setting, requiring that stimulus information be held and compared in short-term working memory. Pinto et al. also analyzed the neural signatures of perceptual decision-making in multi-module RNNs trained using the full-FORCE algorithm (DePasquale et al., 2018; Pinto et al., 2019). Both studies found that modular architectures resulted in substantially different population dynamics than networks without modules. Consistent with this, our findings show that information encoding differs even at the level of single neurons in multi-module networks, a difference that is borne out strikingly in neural recordings from monkeys performing the same task.

One important benefit of artificial neural network models for studying neural systems is the way that they facilitate causal perturbations to test putative circuit mechanisms. In vivo, these perturbations are usually targeted based on features like anatomy or genetic identity. Extending beyond the animal experiments, we adopt an alternative targeting approach, selectively and gradually silencing the activity of artificial units based on the extent to which they show a particular pattern of information encoding—in this case, nonlinear integration of sequentially presented stimuli. Using this type of functional targeting, we validated the necessity of nonlinear integrative encoding for mediating the M/NM decisions during the DMC task. This is an important complement to our experimental results which demonstrate a correlation between the activity of PFC NINs with trial-by-trial decisions. It also demonstrates the utility for RNNs to directly test putative circuit computations underlying cognitive functions, and to motivate future experiments to test model predictions.

It will be important to extend this work to examine the roles played by a wider network of cortical and subcortical areas in solving sequential decision tasks. This includes premotor cortex, which shows decision related activity during a shape (cat vs. dog) DMC task (Cromer et al., 2011) and abstract decision tasks (Wallis and Miller, 2003) as well as subcortical structures such as basal ganglia and thalamus. Furthermore, it will be interesting to characterize multi-module RNNs to explore the local and long-range circuit mechanisms underlying different cognitive functions, that would otherwise be unattainable through current experimental tools.

## ACKNOWLEDGMENTS

We thank Dr. Pantea Moghimi, Krithika Mohan and Barbara Peysakhovich for their constructive and helpful comments during the manuscript preparation.

## AUTHOR CONTRIBUTIONS

SS and DJF designed the main experiments. SS trained monkeys and collected the behavioral and neuronal data for the main experiments. NM and DJF designed the experiment used in Figure 3-figure supplement 1, NM trained monkeys and collected the data for that experiment. YZ, MR and DJF designed the network models, MR trained the network models and implemented the RNN inactivation experiment. YZ analyzed the data and made figures. YZ wrote the manuscript. OZ, MR and NM edited the manuscript. DF supervised the experiments and edited the manuscript.

## GRANTS

This study is supported by NIH R01EY019041, NSF NCS 1631571, and DOD VBFF (DJF).

## COMPETING FINANCIAL INTERESTS

The authors declare no competing financial interests.

## Materials and Methods

### Data sets

This study includes five data sets from two independent experiments. Most of the data are from a DMC experiment that includes a PFC dataset, an LIP dataset and an MIP dataset. Analyses from these datasets have been published previously (Swaminathan and Freedman, 2012; Swaminathan et al., 2013), though unrelated to the present study. The data in Figure 3-figure supplement 1 originated from a DMC learning experiment (Masse et al., 2017).

### Behavioral task and stimulus display

The DMC task has been described previously (Freedman and Assad, 2006; Swaminathan and Freedman, 2012) and is briefly summarized below. In this task, monkeys were trained to release a lever when the categories of sequentially presented sample and test stimuli matched, or hold the lever when the sample and test categories did not match. Stimuli consisted of six motion directions (15°, 75°, 135°, 195°, 255°, 315°) grouped into two categories separated by a learned category boundary oriented at 45° (**Figure. 1b**). Trials were initiated by the monkey holding the lever and keeping central fixation. Monkeys needed to maintain fixation within a 2° radius of a fixation point through the trial. 500 ms after gaze fixation was maintained, a sample stimulus was presented for 650 ms, followed by a 1000 ms delay and a 650 ms test stimulus. If the categories of the sample and test stimuli matched, monkey needed to release a manual touch-bar within the test period to receive a juice reward. Otherwise, monkeys needed to hold the touch-bar during the test period and a second delay (150 ms) period, and wait for the second test stimulus, which was always a match, and then release the touch-bar, so that monkeys concluded all trials with the same motor response (lever release). The motion stimuli were high contrast, 9° diameter, random-dot movies composed of 190 dots per frame that moved at 12°/s with 100% coherence. Task stimuli were displayed on a 21-inch color CRT monitor (1280*1024 resolution, 75 Hz refresh rate, 57 cm viewing distance). Identical stimuli, timing, and rewards were used for both monkeys in all PFC, LIP, and MIP recordings. Monkeys’ eye positions were monitored by an EyeLink 1000 optical eye tracker (SR Research) at a sampling rate of 1 kHz and stored for offline analysis. Stimulus presentation, task events, rewards, and behavioral data acquisition were accomplished using an Intel-based PC equipped with MonkeyLogic software running in MATLAB(Asaad et al., 2013) (http://www.monkeylogic.net).

In the DMC learning experiment, two other monkeys were trained to perform a slightly altered version of the standard DMC task. Identical setups, stimuli, timing, and rewards were used, however, only 24 stimulus conditions (sample-test-direction combinations) were used. Neuronal activity was recorded while monkeys learned this DMC task, which is after the monkeys had learned a delay-match to sample (direction) task.

### Electrophysiological recording

Two male monkeys (Macaca mulatta, 8–10 kg) were implanted with a head post and recording chambers positioned over PPC and PFC. Stereotaxic coordinates for chamber placement were determined from magnetic resonance imaging (MRI) scans obtained before chamber implantation. PFC chambers were centered on the principal sulcus and anterior to the arcuate sulcus, ∼27.0 mm anterior to the intra-aural line. Areas LIP and MIP were accessed from the same PPC chamber, which was positioned over the intraparietal sulcus (IPS) centered ∼3.0 mm posterior to the intra-aural line. All experimental and surgical procedures were in accordance with the University of Chicago Animal Care and Use Committee and National Institutes of Health guidelines. Monkeys were housed in individual cages under a 12-h light/dark cycle. Behavioral training and experimental recordings were conducted during the light portion of the cycle.

LIP and PFC recording sessions were interleaved in each monkey to reduce the influence of timing on the neuronal responses and monkeys’ behavior. In monkey A, 35 PFC recordings sessions were followed by 29 LIP sessions and an additional 15 PFC sessions. In monkey B, most LIP recordings (n = 22 sessions) were conducted first, followed by PFC recordings (n = 36 sessions) and simultaneous LIP-PFC recording sessions (n = 4 sessions). The MIP recordings were conducted in separate sessions after completing PFC and LIP recording.

All recording equipment and procedures were the same as in the previous studies (Swaminathan and Freedman, 2012; Swaminathan et al., 2013). LIP and MIP recordings were conducted using single 75-μm tungsten microelectrodes (FHC), a dura piercing guide tube, and a Kopf (David Kopf Instruments) hydraulic micro-drive system. In general, LIP cells were found at more lateral locations and MIP cells were found at more medial locations within the same recording chamber. LIP was 2-7mm below the surface and MIP was 1-5mm below the surface in both monkeys. PFC recordings were made using 250-μm dura-piercing tungsten microelectrodes (FHC) and a custom manual micro-drive system that allowed simultaneous recordings from up to 16 electrodes. Neurophysiological signals were amplified, digitized and stored for offline spike sorting (Plexon) to verify the quality and stability of neuronal isolations. The offline spike sorting used the same standard as in the previous studies which ensured that each single neuron was well isolated.

In the DMC learning experiment, two additional monkeys (Macaca mulatta, 9–12 kg) were implanted with a head post and two 32-channel semi-chronic recording systems (Gray Matter Research) on PPC and PFC. MRI scans were used to guide chamber placement. For PPC recordings, chambers were placed over the IPS, ∼2.0 mm posterior to the intra-aural line and ∼14.0 mm lateral from the midline for monkey Q, and ∼2.0 mm anterior to the intra-aural line and ∼13.0 mm lateral from the midline for monkey

W. We advanced all PPC electrodes until their estimated positions were below the IPS, guided by its known anatomical depth. Additional evidence for electrode depth on many recording channels was the marked reduction in spiking activity as electrodes entered the sulcus. For PFC recordings, chambers were placed over the principal sulcus, ∼29.0 mm anterior to the intra-aural line and ∼20.0 mm lateral from the midline for monkey Q, and ∼33.0 mm anterior to the intra-aural line and ∼22.0 mm lateral from the midline for monkey W. Each micro-drive system contained 32, 125-μm tungsten microelectrodes (Alpha-Omega). Before each session, we lowered electrodes between 0 and 1 mm to optimally record the spiking activity of well-isolated neurons. Neuronal activity in PFC and PPC were recorded simultaneously for every session. The PPC recording might include both LIP and MIP. We used the same standard for offline isolation of single neuron as in the regular DMC experiment. All experimental and surgical procedures were standard and in accordance with the University of Chicago Animal Care and Use Committee and National Institutes of Health guidelines.

### Receptive field mapping and stimulus placement

All PFC and LIP neurons as well as most MIP neurons were tested with a memory-guided saccade (MGS) task before DMC task. LIP neurons were identified by spatially selective visual responses and/or persistent activity during the MGS task. MIP neurons were identified by responses during the animals’ spontaneous hand movements, such as lever releases, scratching, or arm movements observed before the DMC task commenced, and the absence of modulation during the MGS task. LIP and MIP neurons were also differentiated based on anatomical criteria, such as the location of each electrode track relative to that expected from the MRI scans, the pattern of gray–white matter transitions encountered on each electrode penetration, and the relative depths of each neuron.

Motion stimuli for the DMC task were always targeted to LIP receptive fields (RFs). The typical eccentricity of stimulus placement for LIP recordings was ∼6.0–10.0°. During MIP recordings, the motion stimulus was always placed at 7° from the fixation along the horizontal axis contralateral to the recording hemisphere. For most PFC recordings (n = 55 of 86 sessions), sample and test stimuli were presented in blocks of 30 trials at three non-overlapping locations in the contralateral visual field centered 7.0° from fixation, which covered much of the contralateral visual field on the monitor. For the remaining PFC recording sessions (31 of 86), stimuli were shown at a single fixed location, 7.0° from fixation along the horizontal axis in the contralateral visual field. All recorded trials for PFC neurons were used for subsequent analyses. Similar results were observed using only the one-location or three-location PFC datasets, or using PFC data for which stimuli were presented at the best of the three locations. None of the recorded neurons were pre-screened for direction, category, or M/NM selectivity before recording.

In the DMC learning experiment, we recorded all neurons with well-isolated action potentials, as we could not place stimuli within the RFs of all recorded neurons. To increase the chances that stimuli were in or near neuronal RFs, we ran the experiment in alternating 10-trial blocks in which stimulus position was varied between two non-overlapping positions (7.0° eccentricity; ± 45° relative to horizontal meridian) in the visual field that was contralateral to the hemisphere targeted for neuronal recordings. Analysis of neuronal data revealed qualitatively similar results (in both cortical areas) for each of the two stimulus locations considered separately, or when trials from the two locations were combined. Thus, we combined trials for both stimulus locations in current study.

### Data analysis

#### Pre-analysis neuron screening

We used multi-electrodes to record PFC neurons and did not pre-screen neurons prior to recording. For LIP and MIP recording, we used single electrode recording and applied some standard criteria to screen the neurons (visual responsiveness for LIP and movement responsiveness for MIP). Thus, the total number of neurons was much larger in PFC than in LIP and MIP (PFC: 447; LIP: 75; MIP: 94). However, many PFC neurons exhibited very low firing rates and were not task-modulated during the task interval, and therefore might not contribute to the task variable representations. In contrast, most of the recorded LIP and MIP neurons showed relatively high firing rates and were task modulated. To reduce any potential confounds that might be caused by differences between PFC and PPC (LIP and MIP) datasets, we used the following criteria to further pre-screen all neurons for data analysis: (1), the maximum of the mean conditional averaged firing rate during the task interval (from fixation onset to 350 ms after test onset) should be no less than 5 spikes/s; (2), the activity should exhibit at least one kind of task-related modulation (such as: sample category selectivity, test category selectivity, and M/NM selectivity, one way ANOVA test, p < 0.01) during one of the four task intervals (sample period, earlier delay period, later delay period and test periods). After screening, 145 PFC neurons, 53 LIP neurons, and 66 MIP neurons were included for further analysis. We also tested different thresholds (1 spike/s or 4spike/s) to screen the neurons, which produced similar results.

In order to select neurons that showed significant M/NM selectivity during the test period, we applied a one-way ANOVA test (p < 0.01) to the mean activity within a 200 ms time window, slidding by 5 ms, during the test period (50-350 ms after test stimulus onset). To compare M/NM selectivity time courses across different cortical areas (Figure. 3), we only selected neurons that showed significant M/NM selectivity during the early test period (50-300 ms after test onset). The results were qualitatively similar when we used different time windows (50-250 or 50-350 ms after test onset) to select neurons.

In the DMC learning experiment, we used the same criteria with one difference to screen neurons for the data analysis. Since most neurons exhibited a very low firing rate, we selected the neurons that had a maximum firing rate greater than 4 spike/s during the task interval to include more neurons. We also used different thresholds (1 spike/s or 5spike/s) to screen neurons and obtained similar results.

#### Behavioral performance quantification

For all recording sessions that contained trials for all 36 stimulus conditions, we calculated the monkey’s accuracy for each condition within a single session and then averaged across all sessions. To compare performance across PFC, LIP, and MIP datasets, we first calculated the overall average accuracy for each session and then applied a one-way ANOVA test to test whether there were any differences between different datasets.

We separated both match trials and non-match trials into easier and more-difficult sub-groups based on sample and test motion directions as well as monkey’s averaged performance across all recording sessions (including all PFC, LIP and MIP data) for all the 36 stimulus conditions separately for each monkey. There were 10 stimulus conditions in which the motion direction of either sample or test or both stimuli were center direction (135° or 315°) for both match and non-match trials. We defined 9 of the 10 stimulus conditions in which monkeys showed higher average accuracy for both match and non-match trials as easier sub-group and the other 9 stimulus conditions as more-difficult sub-group. Thus, there were roughly an equal number of trials between the easier and more-difficult sub-groups for each sample and test category. To correlate M/NM selectivity with monkeys’ reaction time (RT), we separated the match trials into faster and slower RT trials (below or above median RT) for all conditions in each session; the faster and slower RT trials from two category conditions were pooled together.

#### Spike density function and normalized activity

For all the figures showing the activity of example neurons and population neurons, we used a 20 ms Gaussian window to smooth the PSTH. In figure 4, the activity of each neuron was normalized by its maximum firing rate.

#### Equating (Decimating) firing rates

We equated the firing rate between match preferring and non-match preferring neurons when their M/NM selectivity was compared. For each brain area, we first computed a ratio (R) of the averaged firing rate of the non-match preferring neurons over the averaged firing rate of match preferring neuron during the test period. Since the averaged firing rate of match preferring neurons is higher than that of non-match preferring neurons in all three cortical areas, for each match preferring neuron, we then randomly removed from respective spike trains a number of action potentials that corresponded to the rounded product of 1-R with the number of action potentials for each trial.

#### Receiver operating characteristic (ROC) analysis

We applied ROC analysis to the distribution of firing rates (50 ms sliding time window with a 5 ms step) of each neuron during the test period to quantify their M/NM selectivity. The area under the ROC curve is a value between 0.0 and 1.0 indicating the performance of an ideal observer in assigning M/NM choice based on each neuron’s trial-by-trial firing rates. Values of 0.0 and 1.0 correspond to strong encoding preference for non-match or match, respectively. Values of 0.5 indicate no M/NM selectivity.

To test whether the M/NM selectivity in PFC (Figure. 7) correlated with monkeys’ trial by trial choice, we used ROC analysis to evaluate the activity change in error-match trials relative to correct-match trials. Since the number of correct trials greatly exceeded error trials and might influence the reliability of ROC values, we applied a shuffling procedure to equalize the trial totals between correct and error trials. We first randomly selected the same amount of correct trials as the error trials, and calculated the ROC value. Then, we repeated this procedure 100 times, and averaged the 100 ROC values. The ROC values were calculated in slightly different ways for match-preferring and non-match preferring neurons: values greater than 0.5, indicate that neuronal activity on incorrect match trials was more similar to activity on correct non-match trials, i.e., lower activity in error-match trials than in correct-non-match trials for match-preferring neurons, or greater activity in error-match trials than in correct-non-match trials for non-match preferring neurons. This is consistent with a correlation between neurons’ M/NM selectivity and monkeys’ trial-by-trial M/NM choices. ROC values near 0.5 indicate similar activity between incorrect match and correct match trials, which indicates that neuronal activity is not correlated with the monkeys’ trial-by-trial choices. While ROC values lower than 0.5, indicate even greater M/NM selectivity between incorrect match and correct non-match trials.

#### Unbiased fraction of explained variance (FEV)

To quantify M/NM and category selectivity, we performed one-way ANOVA on the neuron’s average firing rate within a sliding window (width = 50ms, step size = 5ms), using either the M/NM choice or the category membership as factors. To quantify the amount of information that a neuron encoded about each factor that was independent of the absolute neuronal firing rate, we calculated the unbiased fraction of explained variance (FEV) in the neuron’s firing rate that could be attributed to the M/NM choice or category membership (sample category or test category) with the following:

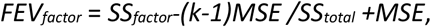

where *SS* indicates the sum of squares, MSE indicates mean square error, and k indicates number of conditions.

We also calculated the unbiased FEV of M/NM choice for the LFP signal. We directly applied the analysis on the average amplitude of the LFP signal within a 50 ms sliding window in all recording channels of each cortical area. The LFP signal was pre-filtered by a band stop filter (Butterworth, 59∼61 HZ) to remove power-line noise.

#### ROC-based category tuning index (rCTI)

We used the rCTI measurement to quantify the category selectivity, which was described in detail in our previous work (Swaminathan et al., 2013) and defined as follows:

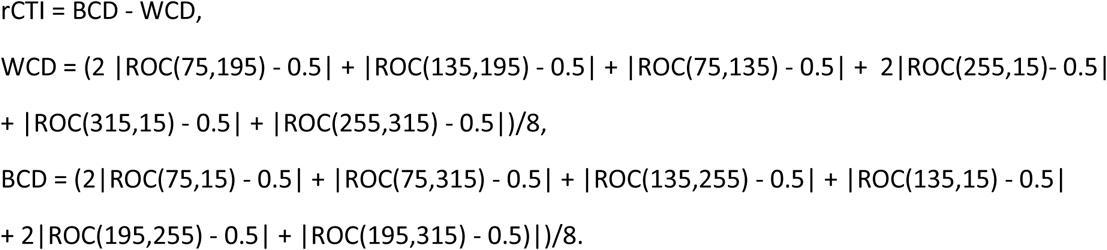

#### Latency of M/NM selectivity

We calculated the latency of M/NM selectivity for spiking activity or LFP signals using the following criteria: 1) the activity (spike count or mean LFP amplitude) in at least two successive (following) sliding time windows (20ms width, 20 ms step size) showed significant M/NM selectivity (one-way ANOVA test, p < 0.05); 2) the M/NM preference of the activity in these time windows must be consistent with the global M/NM preference during the test period (50∼350 ms after test onset). For this calculation, we only analyzed neurons that showed significant M/NM selectivity. When we compared the latency between different sub-groups or brain regions, we used only the neurons that we could calculate a latency according to these criteria. We also tested different criteria (different numbers of sliding window such as 5) to perform the analysis and obtained similar results.

#### Correlation between sample category and test category representation and selection of neurons showing mixed category selectivity

To test the correlation between sample and test category selectivity during the test period, we performed one-way ANOVA on the neuron’s activity and calculated the unbiased FEV of both sample category and test category as mentioned above during the early test period (0-250 ms after test stimuli onset), which mostly preceded the monkeys’ decision. The analysis was applied on the mean firing rates of each neuron with a 100 ms sliding window (5 ms step). The maximal FEV value of sample category and the maximal FEV value of test category of each single neuron were chosen to calculate the rank correlation between sample and test category selectivity in each cortical area. Neurons that showed significant selectivity for both sample and test category in at least one of the selected sliding windows above (0∼250 ms after test onset, p < 0.01) were defined to be mixed-category-selective neurons.

#### Support vector machine (SVM) decoding

Similar to previous studies (Sarma et al., 2016), we used a linear SVM classifier to decode monkeys’ M/NM choice, category membership, and sample-test-category combination separately from a surrogate population of three cortical areas. In this surrogate population, activities from different neurons in one cortical area were treated as if they were recorded simultaneously although neurons were, for the most part, not recorded simultaneously. The linear classifier was trained using an SVM. In training a linear classifier, a hyperplane that best separates the trials belonging to two or several different classes was determined. In the case of decoding M/NM choice, each class corresponds to one type of choice. In contrast to previous studies (Swaminathan et al., 2013), we used all six motion directions together to perform the decoding analysis for category, as we think the direction turning might also contribute to the category representation.

Decoding was applied to the mean firing rates of neurons within a 50 ms sliding window (5 ms step). For each neuron, we randomly selected 66% of trials to train the classifier and left the other 34% of trials for testing. We then randomly sampled, with replacement, 120 trials from the training list and 60 trials for the testing list for bootstrapping. In order to reduce the potential confound caused by uneven number of trials of different motion directions, a minimum number for trials of each motion direction was required for random sampling (10 and 5 trials for training and testing data of each direction, respectively). To compare different types of selectivity across different populations, we applied a shuffling procedure to select an equal number of neurons for all decoding analysis except in Figure. 4. We bootstrapped all decoding analyses 100 times.

#### Nonlinearity index

To quantify the nonlinearity of mixed sample and test category selectivity, a nonlinearity index was calculated for each mixed category selective neuron in PFC and LIP. The nonlinearity index was defined as the test category selectivity difference between two sample category conditions:

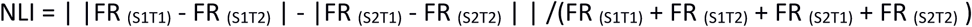

where NLI indicates nonlinearity index, ‘| |’ indicates absolute value, and FR indicates mean firing rate. On one hand, if the test category selectivity is linear addictive to the existing sample category selectivity, then (S_1_T_1_ - S_1_T_2_) = (S_2_T_1_ -S_2_T_2_). On the other hand, if the neuron purely responds to the M/NM status of the test stimuli, then (S_1_T_1_ - S_1_T_2_) = -(S_2_T_1_ -S_2_T_2_). Their absolute value would be the same.

Therefore, if there is pure linear mixed sample-test-category selectivity or pure M/NM selectivity, the value of nonlinearity index will be 0.

#### Recurrent neural network training

All RNN analyses involved training biologically-inspired networks, as described previously (Masse et al., 2019). These differ from standard RNNs in two main ways: first, they contain separate excitatory and inhibitory units, per Dale’s Law; and second, synapses are endowed with short-term plasticity, allowing synaptic efficacies to fluctuate over short timescales in an activity-dependent manner, as in a previous study from our group. All networks were trained using Tensorflow (Martín Abadi et al., 2016) on a GEFORCE RTX-2080Ti GPU with the same hyperparameters: 400 hidden units, 320 of which were excitatory and 80 of which were inhibitory; a learning rate of 0.002; a batch size of 512 trials; metabolic costs on mean activity and mean connection strength, weighted consistently across all networks; and initial weights and biases for input, hidden, and output weight matrices drawn from identical distributions. All networks received input from 24 motion-direction-tuned neurons, with tuning distributed according to a von Mises function with concentration factor 2 and a scaling factor of 4. All networks were wired to 2 output units, which corresponded with the animals’ behavior of holding/releasing a touch-bar to indicate its match/non-match decision on each trial.

We trained all 50 networks to perform three benchmark cognitive tasks: delayed match-to-sample (DMS), delayed match-to-rotated-sample (DMRS), and delayed match-to-category (DMC) (Masse et al., 2019). This was motivated by the observation that networks trained to perform DMC alone often adopt idiosyncratic solutions to the task, where categorical tuning is immediately established by adjusting input weights and direction information is immediately discarded. To our knowledge, this kind of fine tuning at the level of sensory inputs to higher cortical areas is not consistent with neurophysiological results, as both category and direction information was encoded in LIP during the sample period of the DMC task. We thus chose to interleave training on DMC with training on DMS and DMRS–both of which require direction information to be preserved–to encourage more biological network solutions. All networks achieved consistently high performance on all three tasks by the end of training (>95 percent accuracy for the last 50 batches).

#### Implementing multi-module RNNs

The existence of multiple modules in these RNNs was implemented through constraints on the initial recurrent connectivity of the hidden layer. The simplicity (and computational efficiency) of this approach for implementing multi-module RNNs derives from the way that the separation between excitatory and inhibitory units is implemented: all connection weights are passed through a ReLU before they are multiplied by a constant +1/-1 (E vs. I) and applied, so a connection that is culled before training never contributes to the loss, and is never adjusted up or down. To model LIP and PFC, we built networks with two modules, with half of the hidden layer units designated to each module. Each module was allocated half of the excitatory units and half of the inhibitory units in the overall network to ensure that the modules did not differ in their balance of excitation/inhibition prior to training. Motivated by the observation that inhibitory connections in cortex are largely local, we prohibited all inhibitory projections targeting out-of-module neurons. Divisions between brain areas are also distinguished by denser connectivity within areas than between areas, a form of bottleneck that we modeled by restricting the number of excitatory connections between modules to at most 10 percent (40 units) of their possible volume. We implemented a similar bottleneck on connections with respect to sensory inputs and motor outputs. Only 10 percent of units (40) in module 1 (LIP-like module) could receive projections from the motion-tuned input units; similarly, only 10 percent of units (40) in module 2 (PFC-like module) could project to the output neurons. To encourage the functional differentiation of these modules, we prohibited out-of-module projections from neurons receiving motion-tuned inputs/projecting to motor outputs.

#### Analysis of RNN activity

We performed the same analyses on units in the RNNs as we did on the neurophysiology data. As with the neural recordings, we only included the units which showed task-related activity during the test period of the DMC task, defined using the following criteria: (1) the maximum of the averaged activity during the test period should be no less than 0.001; (2) the activity should exhibit at least one kind of task-related modulation during the test period (such as: sample category selectivity, test category selectivity, and M/NM selectivity, one way ANOVA test, p < 0.01).

#### Inactivation experiments in silico

To assess the contribution of different RNN units to M/NM decisions, we performed an in silico analogue of neuronal inactivation experiments similar to those used in experimental studies. The monkey experiments revealed a correlation between the activity of PFC nonlinear integrative neurons and the animals’ trial-by-trial decisions. As such, we hypothesized that inactivating units with greater nonlinear integrative encoding would have a greater impact on the RNN’s ability to perform the task than inactivating units with weaker nonlinear integrative encoding. To test this hypothesis, we examined the RNNs’ behavioral performance after inactivating different subsets of units in the PFC for the duration of the test period during the DMC task (the final 650ms of each trial). To do this, we first freeze all the parameters of the RNNs after the initial learning. We then divide task-related units in the PFC module into more-nonlinear and less-nonlinear groups, as measured by nonlinearity index. The more- and less-nonlinear groups might project to the output units to different extents, a difference which could explain any divergence in network behavior during inactivation across groups rather than the specific ablation of a local network computation. To control for this possibility, we included only those units in the PFC module that did not directly project to the output units in the inactivation analyses. The mean activity of RNN units during the test period varied across a wide range (from 0 to more than 20). This range could produce a bias toward high values of the nonlinearity index for units with very low activity. Such a bias could yield different overall levels of mean network inactivation for different groups of units selected by values of the nonlinearity index. To guard against this confound, we computed the receiver operating characteristic (ROC) value to quantify the overlap between the distributions of firing rates for the two categories in order to compute nonlinearity index. After selecting the two inactivation groups based on these nonlinearity indices, and ensuring that they matched in size, we performed bulk inactivation from the onset of the test stimulus to the end of the trial during DMC. The inactivation was implemented by directly multiplying an activity multiplier (<= 1) to the activity of the target units. In this way, we performed precise, graded causal manipulations -- e.g. to smoothly titrate the amount of inactivation applied to each unit from complete (100% reduction in activity) to minimal (0% reduction in activity).

---

**Figure 2 Supplement 1.**
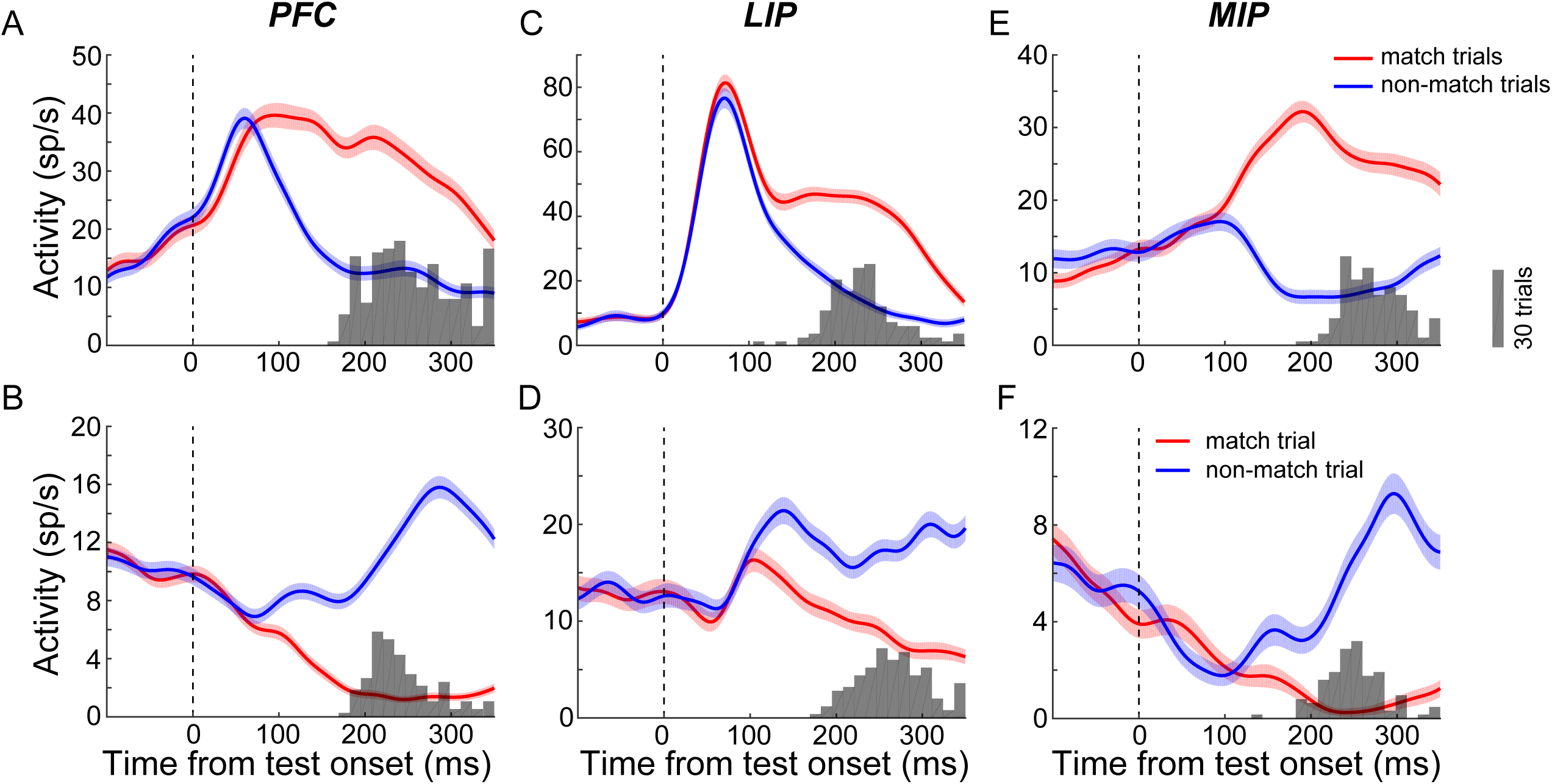
Examples of match-preferring and non-match-preferring neurons in PFC, LIP and MIP. **A**. The averaged activity of a match-preferring PFC neurons during match trial (red) and non-match trial (blue). Shaded area denotes ±SEM. The gray histogram represents the reaction time distribution on match trials. **B**. Activity of a non-match preferring PFC neuron. **C-D**. The activity of match-preferring (C) and non-match-preferring (D) example neurons in LIP. **E-F**. The activity of match preferring (E) and non-match-preferring (F) example neurons in MIP.

**Figure 2 Supplement 2.**
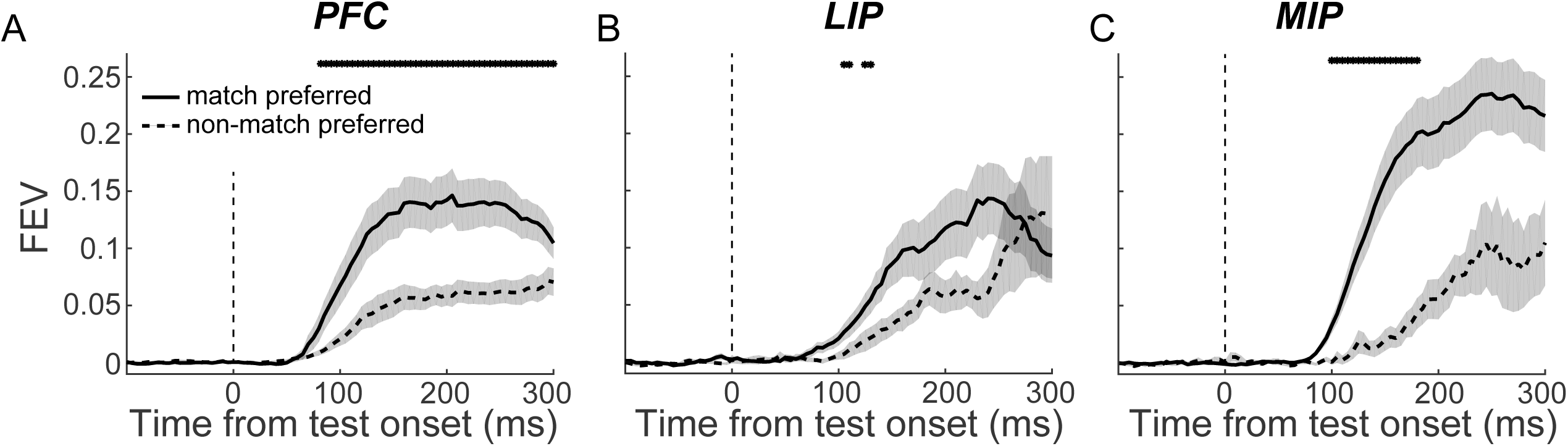
The comparison of the M/NM selectivity between match-preferring and non-match-preferring neurons. **A**. The M/NM selectivity of match-preferring (solid) and non-match-preferring (dashed) neurons in PFC was evaluated using the unbiased FEV. Shaded area denotes ±SEM. **B-C**. The M/NM selectivity of neurons in LIP (B) and MIP (C).

**Figure 3 Supplement 1.**
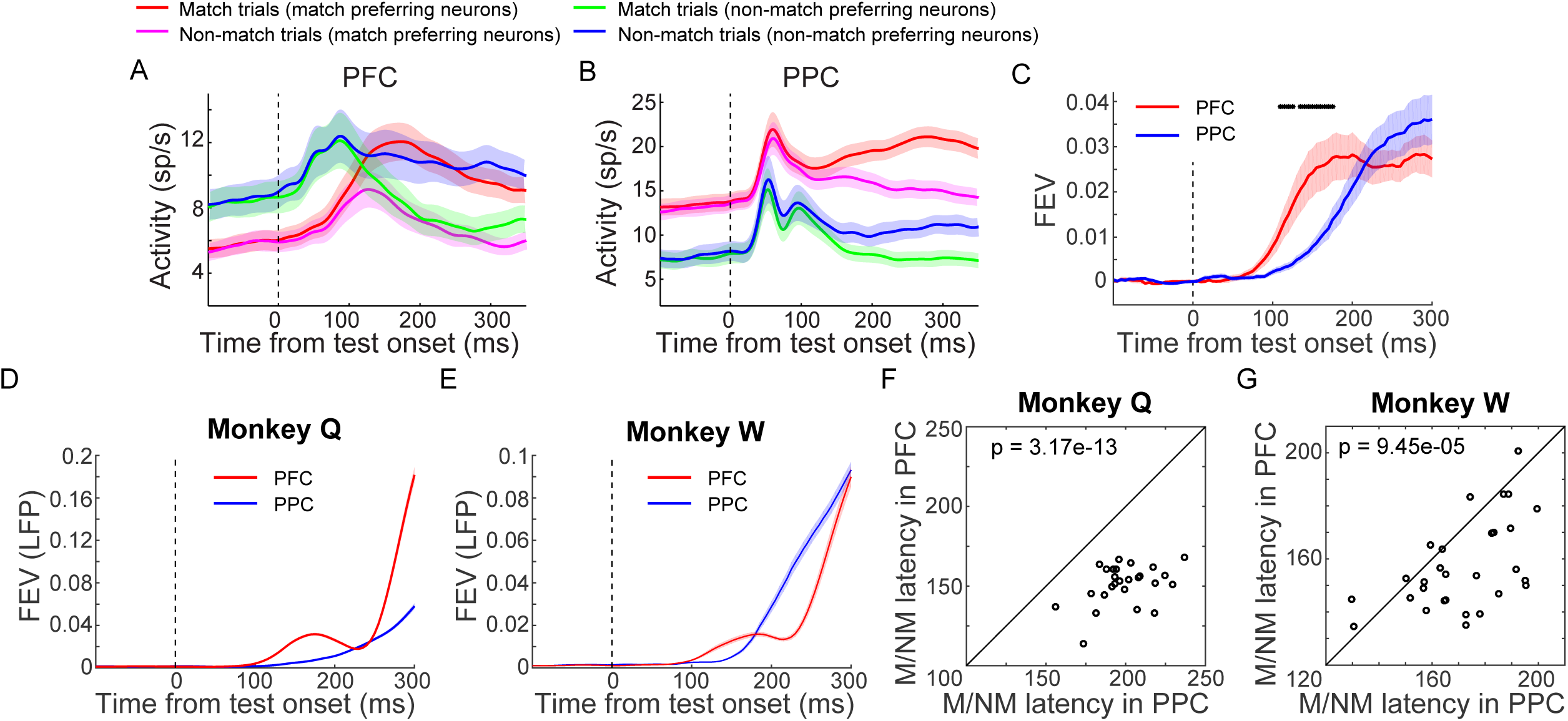
The comparison of M/NM selectivity between PFC and PPC in a DMC learning task. **A-B**. The population activity of match-preferring and non-match-preferring neurons in PFC (A) and PPC (B) are shown as a function of time. **C**. The M/NM selectivity of all PFC (red) and PPC (blue) neurons are evaluated using unbiased FEV. PFC showed significantly earlier M/NM selectivity (P = 0.0158, Wilcoxon test). **D-E**. The M/NM selectivity of LFP amplitude in PFC (red) and PPC (blue) are shown separately for two monkeys. LFP signals were recorded simutaneurouly from PFC and PPC by using multi-channel recording. **F-G**. The comparisons of the mean time point in which LFP signal started to show significantly M/NM selectivity between PFC and PPC (P < 0.01, one-way ANOVA). Each symbol represents the averaged start time in all recording channels within one recording session. The shaded area represents ±SEM.

**Figure 5 Supplement 1.**
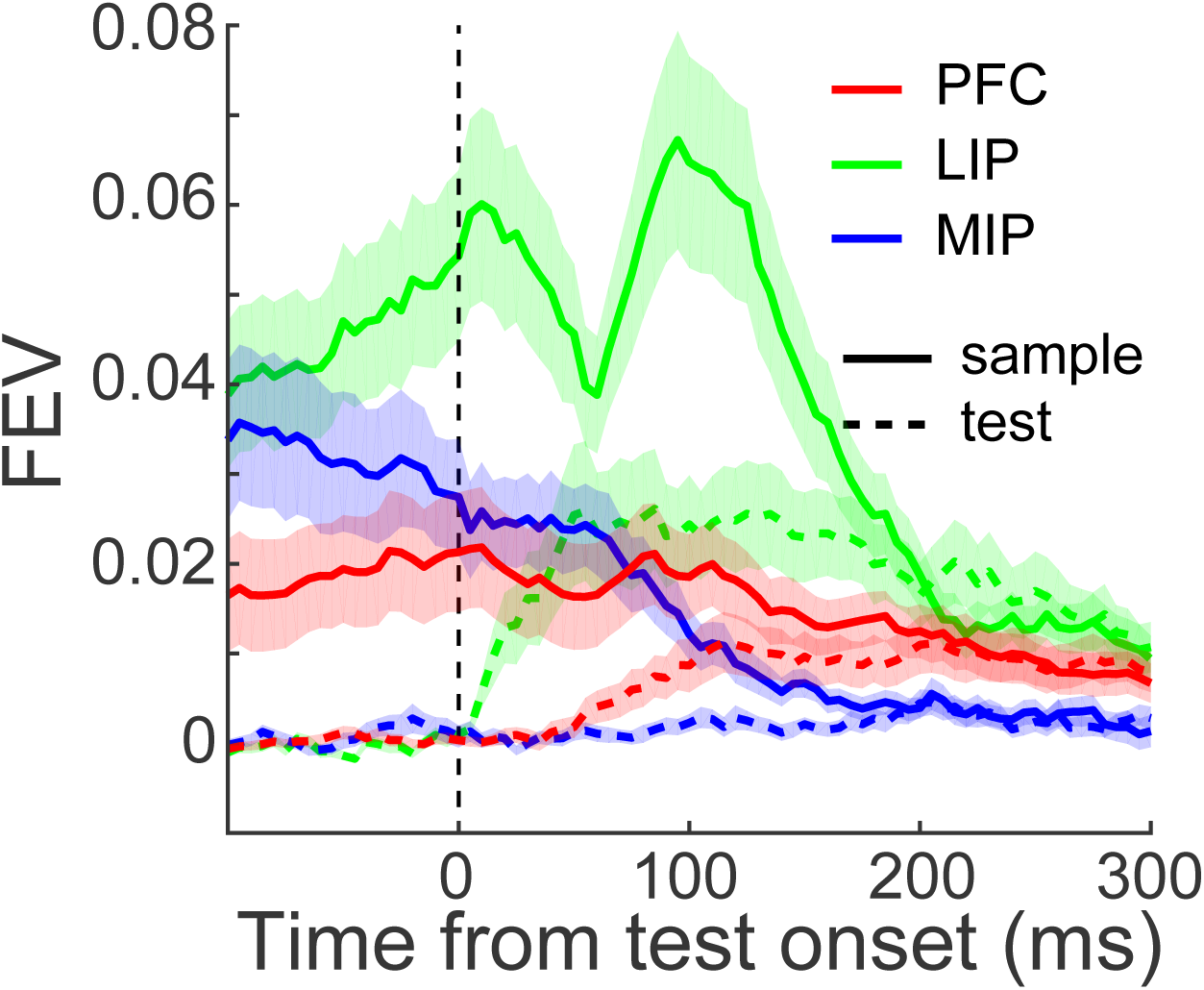
The comparisons of category selectivity among PFC, LIP and MIP. Both sample (solid) and test (dashed) category selectivity of PFC (red), LIP (green) and MIP (blue) neurons were evaluated using unbiased FEV. The shaded area represents ±SEM.

**Figure 8 Supplement 1.**
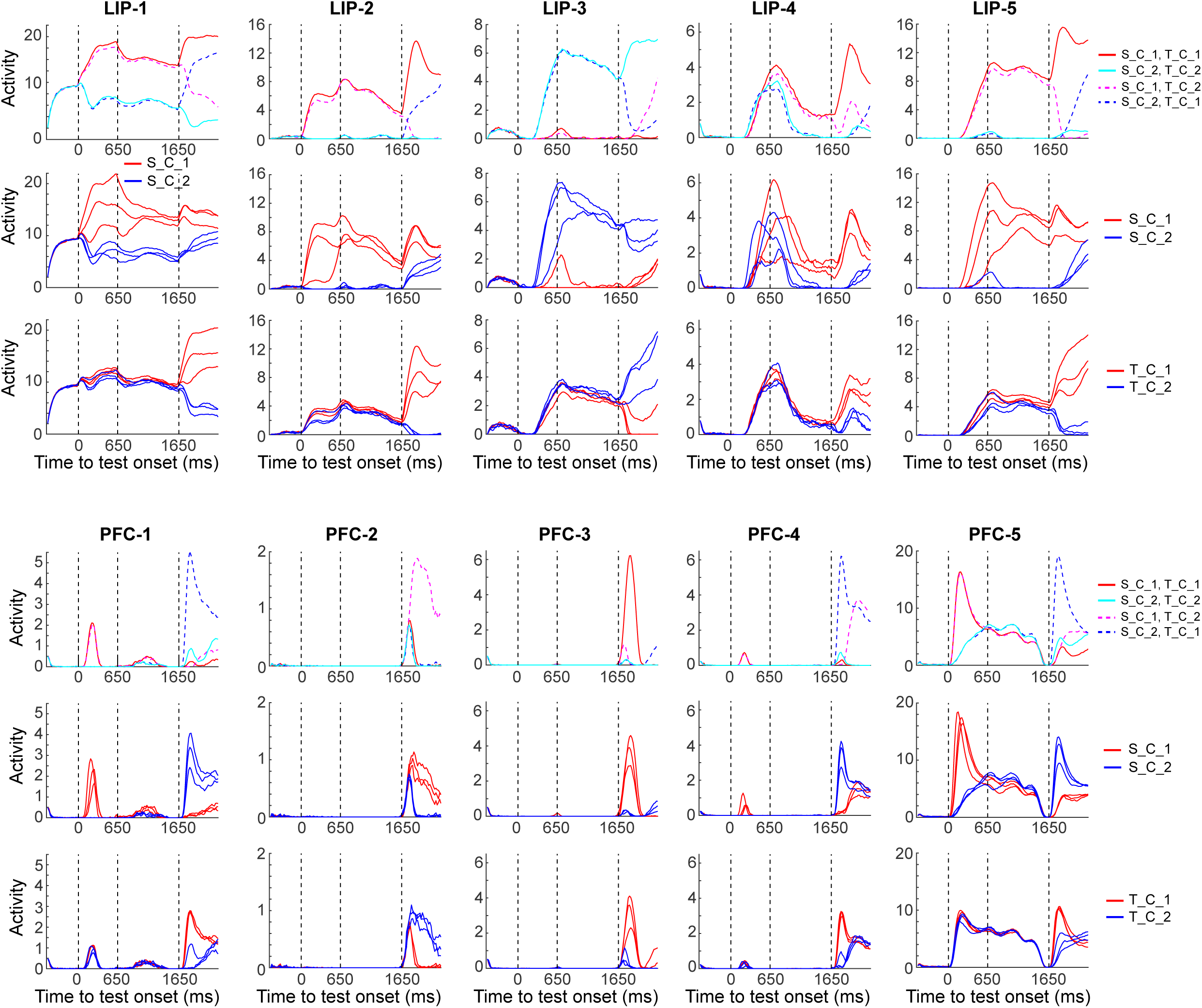
Examples of units showed both sample and test category selectivities during test period of DMC task in both LIP and PFC modules of one example RNN. The upper three rows: five example units form LIP module with each column corresponding to one unit. The first row shows the condition averaged activity sorted by sample and test category combinations. Within each panel, the first and second dashed vertical lines denote the time interval for sample stimulus, while the third dashed vertical line indicates the time of test stimulus onset. The averaged activity for each sample and test directions is plotted in the second and third rows, respectively. Different colors represent different sample category or test categories. The lower three rows show five example units from PFC module.

**Supplementary Table 1:**
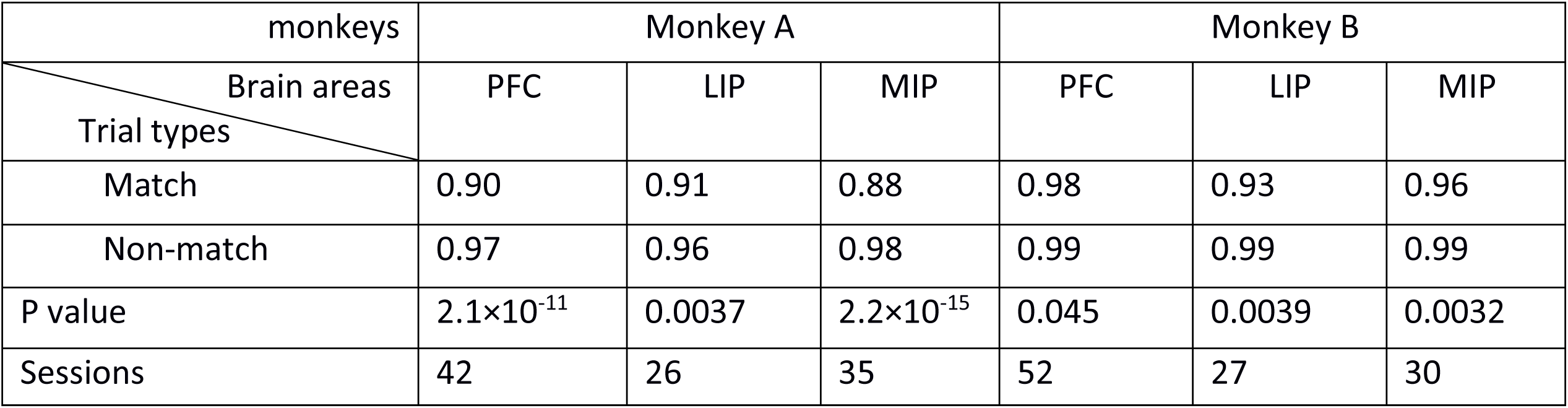
Two monkeys’ M/NM choice accuracy during match and non-match trials.

